# Interactions between the medial prefrontal cortex, dorsomedial striatum, and dorsal hippocampus that support rat category learning

**DOI:** 10.1101/2025.04.21.649785

**Authors:** Matthew B. Broschard, Jangjin Kim, Hunter E. Halverson, Sean J. Farley, John H. Freeman

## Abstract

Categorization creates memory representations that are efficient, generalizable, and robust to noise. Multiple brain regions have been implicated in categorization, including the prefrontal cortex, striatum, and hippocampus; however, few studies have examined how these regions interact during category learning. We recorded neural activity in the medial prefrontal cortex (PFC), dorsomedial striatum (DMS), and dorsal hippocampus (HPC) while rats learned to categorize distributions of visual stimuli. We found a learning-related shift in contributions from the PFC (with DMS→PFC theta (4-10Hz) interactions) to the HPC (with HPC→DMS and bidirectional PFC-HPC theta interactions). Decision-making depended on DMS and HPC spiking, as well as the PFC→HPC→DMS pathway. Our results provide a framework that characterizes how the PFC-DMS-HPC network interacts during category learning. This is informative for multiple neurological disorders that affect category learning, including Parkinson’s Disease, autism, and dementia.

**Highlights:**

- Rats learned to categorize distributions of visual stimuli.
- Early training sessions relied on PFC spiking and DMS→PFC theta interactions.
- After learning, contributions shifted to HPC spiking, HPC→DMS interactions, and bidirectional PFC-HPC interactions.
- Decision-making depended on DMS and HPC spiking and PFC→HPC→DMS interactions.

**Graphical Abstract:** 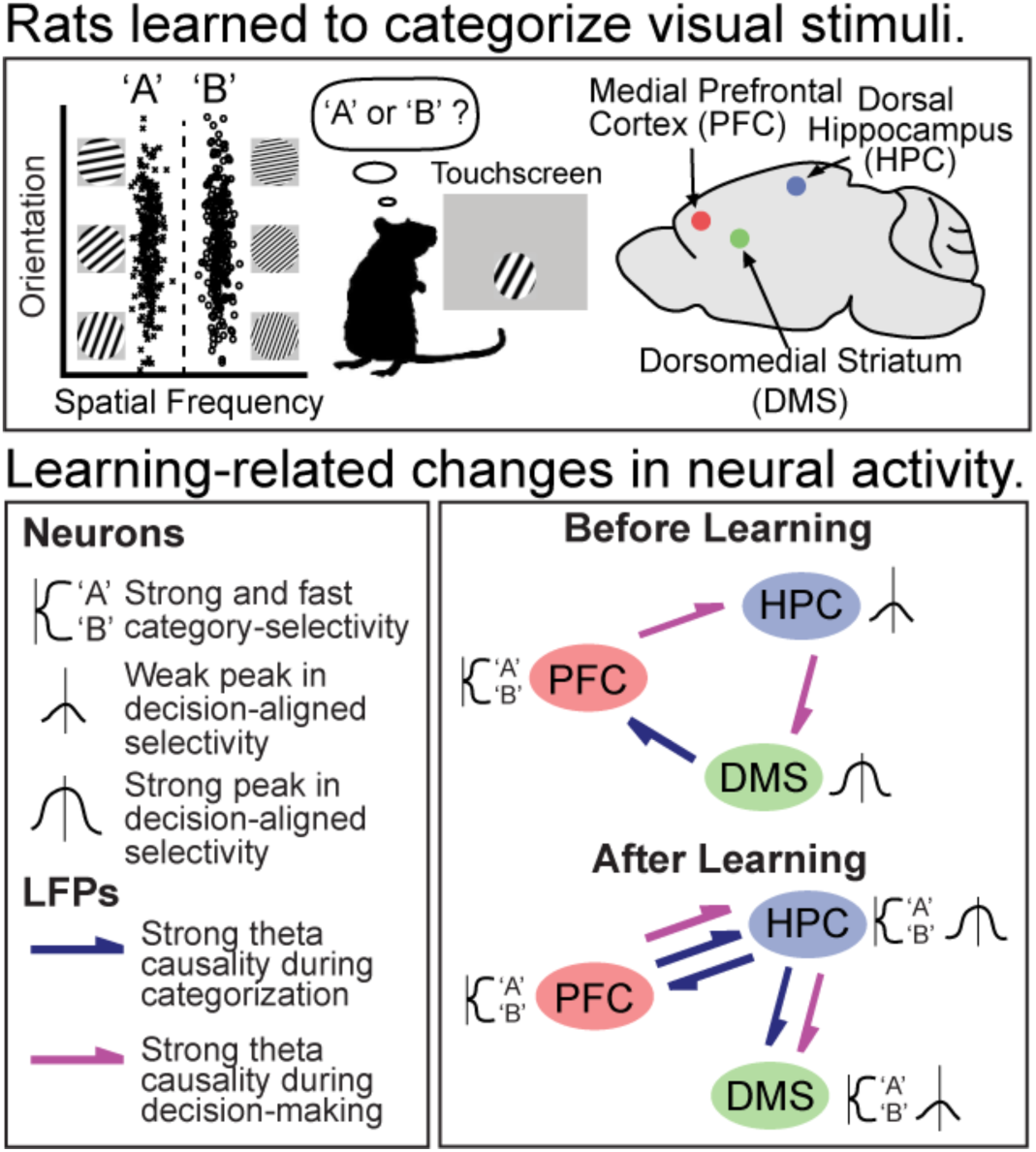

## Introduction

Adaptive behavior requires generalizing knowledge to new situations. This is not trivial, considering our environment contains large amounts of variability. No two events are identical. The brain manages this variability by organizing experiences into categories^1^. Categorization extracts regularities across experiences while ignoring irrelevant details. This allows for memory representations that are robust to noise and can be easily generalizable.

Experiments from the last two decades have identified multiple brain regions that are critical for categorization in humans and monkeys^2–3^. These regions include (but are not limited to) the prefrontal cortex, hippocampus, and the striatum. The prefrontal cortex is thought to use executive functions to identify category-relevant stimulus information and evaluate category rules^4–6^. The hippocampus is thought to build and maintain abstract memory representations^7–9^. The striatum is thought to rapidly associate category stimuli to behavioral responses^10–14^.

Computational models of categorization emphasize the importance of inter-regional interactions during category learning. Traditional models posit that the prefrontal cortex and the striatum form working memory loops that evaluate category rules^14–15^. Newer models posit that interactions between the prefrontal cortex and the hippocampus create flexible, abstract category representations that emphasize category-relevant stimulus information^16–17^. Few studies have tested these predictions directly, and inter-regional communication during category learning is largely unknown.

Our group has found that rat analogs of these regions are also critical for category learning. These include the medial prefrontal cortex (PFC)^18^, the dorsal hippocampus (HPC)^19^, and the dorsomedial striatum (DMS)^20^. Simulations using a network model^21^ showed that these regions were functionally similar to humans. The PFC was correlated with the network’s attention mechanism^18^, the HPC was correlated with the network’s memory representations^19^, and the DMS was correlated with the connection between the memory representations and the category labels^20^. This suggests there is a general correspondence between rats and humans.

In the current experiment, we recorded spiking activity and local field potentials (LFPs) in the PFC, DMS, and HPC simultaneously in rats during category learning. The stimuli contained black and white gratings that varied in their spatial frequency and orientation (Fig. 1A). Only one dimension was required to categorize the stimuli, and the other dimension could be ignored. These stimuli and task procedures are highly similar to those used in human experiments^22^. Analyses focused on examining changes in the spiking activity and network synchrony that characterized how PFC-DMS-HPC interactions developed with learning.

**Figure 1.**
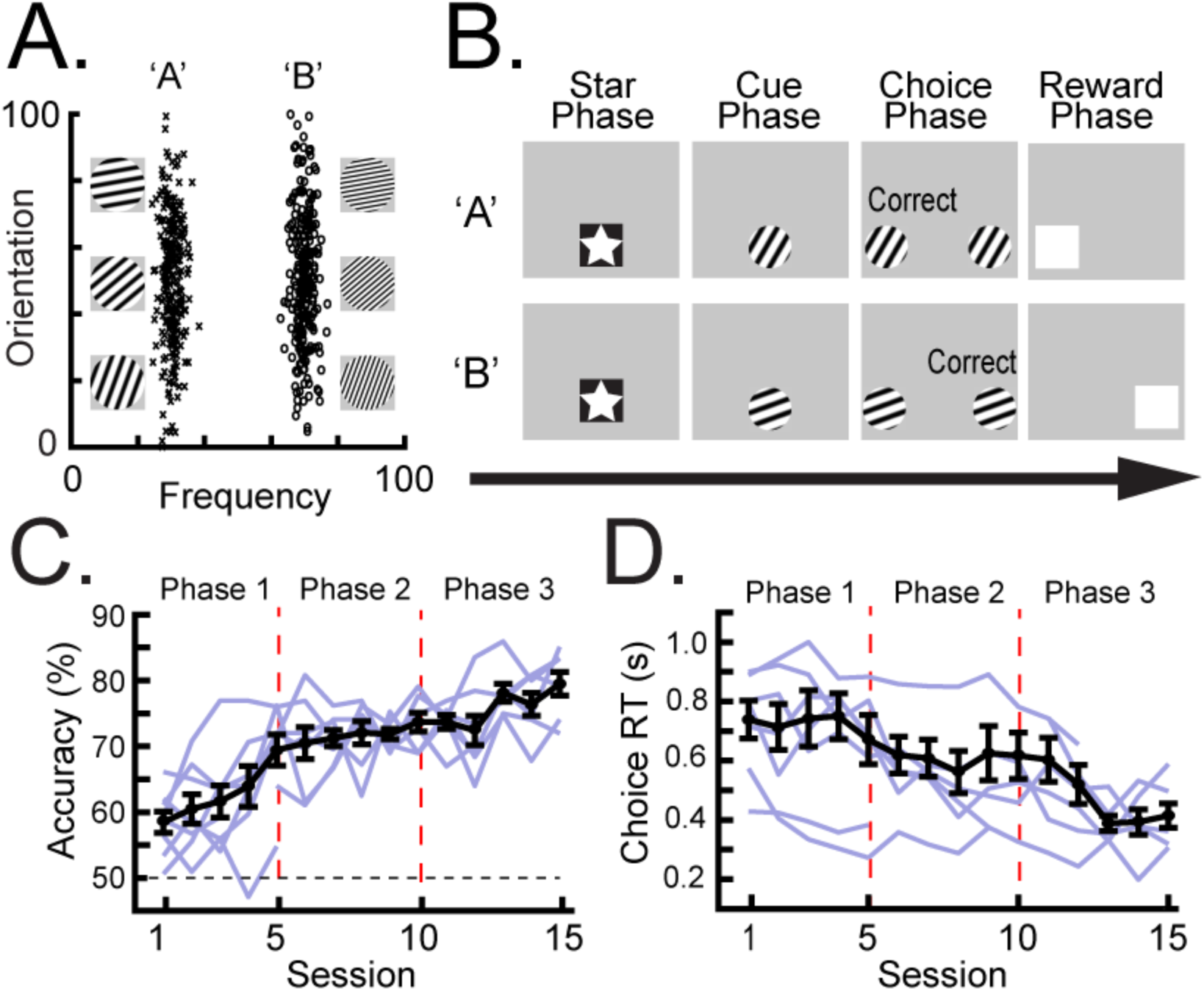
Category learning paradigm. **A,** Rats were trained to categorize distributions of visual stimuli containing black and white gratings. These gratings varied in their spatial frequency and orientation. This task is typically learned by attending to the spatial frequency dimension and ignoring the orientation dimension. **B,** Trial procedure. <u>Star Phase:</u> Each trial was initiated by touching the star stimulus in the center of the screen. <u>Cue Phase:</u> A unique category stimulus was sampled from the category distributions and presented on the screen. The rats had two seconds to touch the category stimulus. <u>Choice Phase:</u> Two report keys appeared on the left and right sides of the screen. The rats touched one of the report keys, depending on the category membership of stimulus presented during the Cue phase (e.g., the left report key for category ‘A’ stimuli and the right report key for category ‘B’ stimuli). <u>Reward Phase:</u> After a correct response, a white box appeared on the screen, and a food pellet was delivered. After an incorrect response, the trial aborted without reinforcement. **C-D,** Training was divided into three phases (i.e., Phase 1, Phase 2, and Phase 3), and each phase averaged data from five training sessions. Background plots indicate individual rats. **C,** Session accuracy increased between each training phase. **D,** Choice RT decreased between each training phase. All error bars indicate *S.E.M*.

## Results

### Training Results

Long Evans rats (n = 9, 3 females) were trained to categorize distributions of visual stimuli containing black and white gratings (Fig. 1A). The rats were shown unique stimuli on every trial (Fig. 1B; Cue phase). They decided the category membership of each stimulus by touching one of two report keys appearing on the left and right sides of the screen (Choice phase; e.g., the left report key for category ‘A’ and the right key for category ‘B’). A pellet reward was delivered after correct responses to guide learning (Reward phase).

All analyses were separated into three training phases (i.e., Phase 1, Phase 2, and Phase 3). Each phase averaged data across five training sessions, which roughly separated the different phases of learning (e.g., before learning, after learning, and retention). Accuracy improved after each training phase (Fig. 1C; *t*(4.56) = 4.40, *p* = .009), and reaction time became faster after each training phase (Fig. 1D; *t*(4.50) = 3.21, *p* = .027). There were no significant differences in performance between male and female rats (accuracy and reaction time, *p* > .05).

### Histology

Spiking activity and LFPs were recorded in the PFC, DMS, and HPC during each session. The PFC tetrodes spanned dorsal portions of the prelimbic area (PL) as well as ventral portions of the rostral anterior cingulate (ACC; Supplemental Fig. 1A). DMS tetrodes were concentrated in the dorsal portion of the DMS. HPC tetrodes were concentrated in the CA1, and a few tetrodes extended into the dentate gyrus (DG). In total, 2,391 separable neurons (PFC: 1203; DMS: 618; HPC: 570) were analyzed (Supplemental Fig. 1A; see Supplemental Fig. 1B for representative examples). Firing rates were similar for neurons in the two PFC subregions (PL vs. ACC; Supplemental Fig. 2A), the two HPC subregions (CA1 vs. DG; Supplemental Fig. 2B), and between male and female rats (Supplemental Fig. 2C). There were an insufficient number of neurons to examine these comparisons across training phases.

### Learning-related changes in neuronal category-selectivity

Neurons showed reliable firing rate differences between category ‘A’ trials and category ‘B’ trials (i.e., “category-selectivity”). To visualize this category-selectivity across neurons, we organized each neuron’s activity depending on the category that elicited a greater response (Fig. 2A; “Higher” vs. “Lower”). In each region, category-selectivity appeared early in the Cue phase and persisted through the Reward phase. This was consistent for neurons with firing rates that increased or decreased from baseline during the trial events (Fig. 2B; Supplemental Figs. 3A-C). Most DMS and HPC neurons had activity that decreased from baseline, whereas the PFC had a more balanced number of excitatory and inhibitory neurons (Supplemental Fig. 3D).

**Figure 2.**
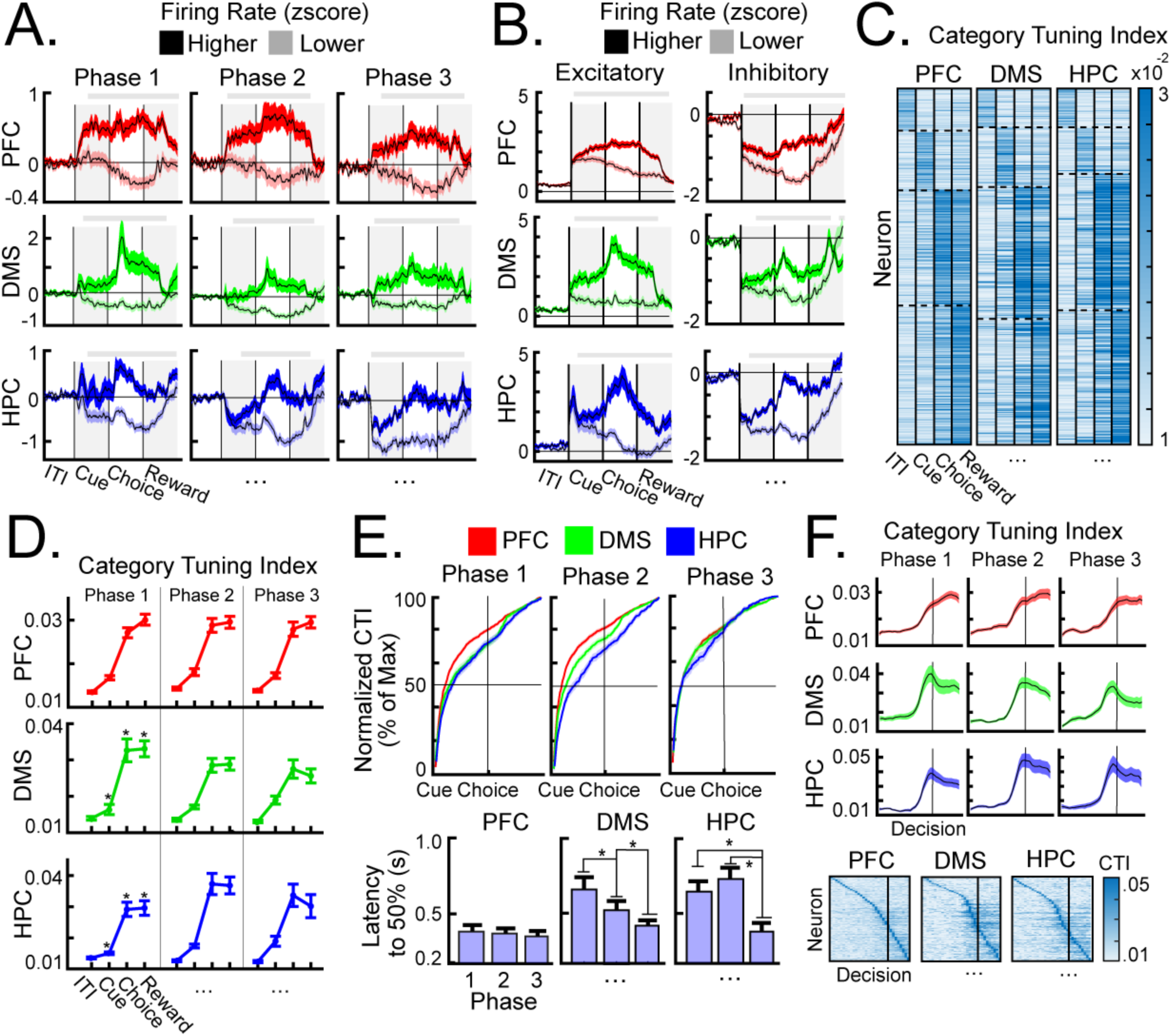
Learning-related changes in neuronal category-selectivity. **A,** Average firing rates, separated by each neuron’s ‘Higher’ and ‘Lower’ categories, for each region and training phase. **B,** Average firing rates for neurons with excitatory activity (left) and inhibitory activity (right), separated by each neuron’s ‘Higher’ and ‘Lower’ categories. **C,** A category tuning index (CTI) quantified category-selectivity at each trial event. Heatmaps of CTI values for each neuron (rows) and trial event (columns). Neurons were sorted according to their maximum CTI value. **D,** During the Cue phase, CTI increased across training phases in the DMS and HPC. During the Choice phase and Reward phase, CTI increased across training phases in the HPC and decreased across training phases in the DMS. There were no significant changes in CTI in the PFC in any trial event. **E,** Top: the speed on selectivity onset, independent of its magnitude, was calculated by normalizing CTI values to each neurons’ maximum value. Bottom: the latency to reach 50% of this maximum. PFC neurons had fast-latency selectivity across all training phases and was faster than the other regions during Phases 1 and 2. Latency in the DMS and HPC became faster across training phases. **F,** Top: CTI values aligned to the rats’ decision during the Choice phase. Average CTI in the DMS and HPC peaked before the rats’ decision. Bottom: Heatmaps of CTI values aligned to the rats’ decision. A large proportion of DMS and HPC neurons had CTI values that peaked around the rats’ decision. All error bars indicate *S.E.M.* Gray bars above each plot indicate timepoints in which firing rates were significantly different between categories.

We quantified category-selectivity at each trial event using a category tuning index^4^ (CTI; Fig. 2C). This index compared firing rate differences for between-category trial pairs (e.g., a category ‘A’ trial and a category ‘B’ trial) and within-category trial pairs (e.g., two category ‘A’ trials). CTI values of the vast majority of neurons increased from baseline and peaked during the Cue phase (∼15% of neurons), the Choice phase (∼35%), or the Reward phase (∼35%; Fig. 2C). Only about 30% of neurons had CTI values that were *significantly* larger than baseline (Supplemental Fig. 4). This is consistent with non-human primate studies^4^ and suggests that many neurons only showed partial increases in category-selectivity.

Category-selectivity increased with learning in the DMS and the HPC and was strong in the PFC throughout training. Average CTI values during the Cue phase increased across training phases in the DMS and the HPC (Fig. 2D; *p <* .001) but did not change in the PFC (*p* = .245). This suggests that the DMS and the HPC became more important to categorization after learning. We calculated the speed of selectivity onset, independent of its magnitude, by normalizing CTI values to each neurons’ peak value (see Methods; Fig. 2E). Selectivity onset was faster in the PFC than the other regions during training Phases 1 and 2 (PFC vs. DMS: *p* = .009; PFC vs. HPC: *p* < .001). Selectivity onset became faster across training phases in the DMS and the HPC (all *p*s < .001) and did not change in the PFC (*p* = .326). Together, this suggests that category representations in the DMS and the HPC became stronger with learning. PFC neurons had fast-latency selectivity onset throughout training, emphasizing its importance during early training sessions.

The DMS and the HPC were also critical for decision-making. We isolated activity related to decision-making by aligning trials to the rats’ touch during the Choice phase. For the DMS and the HPC, firing rate (Supplemental Fig. 5) and CTI (Fig. 2F) peaked around the rats’ decisions. In the PFC, activity continued to increase after the rats’ decisions, without a clear peak (Supplemental Fig. 5 & Fig. 2F). Average CTI values during the Choice phase *increased* in the HPC from training Phase 1 to Phases 2 and 3 (Fig. 2D; *p*s < .01) and *decreased* in the DMS from training Phase 1 to Phases 2 and 3 (*p*s < .01). There were no changes in the PFC (*p*s > .05). These results suggest that the DMS and the HPC were important for decision-making. These contributions shifted from the DMS to the HPC after learning.

The DMS and HPC learned to inhibit irrelevant stimulus information. We examined the representation of the irrelevant stimulus dimension (i.e., orientation) by comparing neural responses to stimuli at the “Center” of the category distributions and stimuli at the “Tails” of the distributions (Fig. 3A). About 60% of neurons in each region had firing rates that peaked (either minimum or maximum) for the Center stimuli (“Prototype” neurons; Fig. 3B). Figure 3C shows the average firing rates of these Prototype neurons. In all regions, selectivity for the Center stimuli was strong during training Phase 1 (Fig. 3D). Selectivity decreased across training phases in the DMS and the HPC (*p*s < .001) but persisted across training in the PFC (*p* = .109). These results indicate that the DMS and the HPC learned to filter out information from the irrelevant stimulus dimension. By contrast, the PFC encoded the irrelevant dimension throughout training.

**Figure 3.**
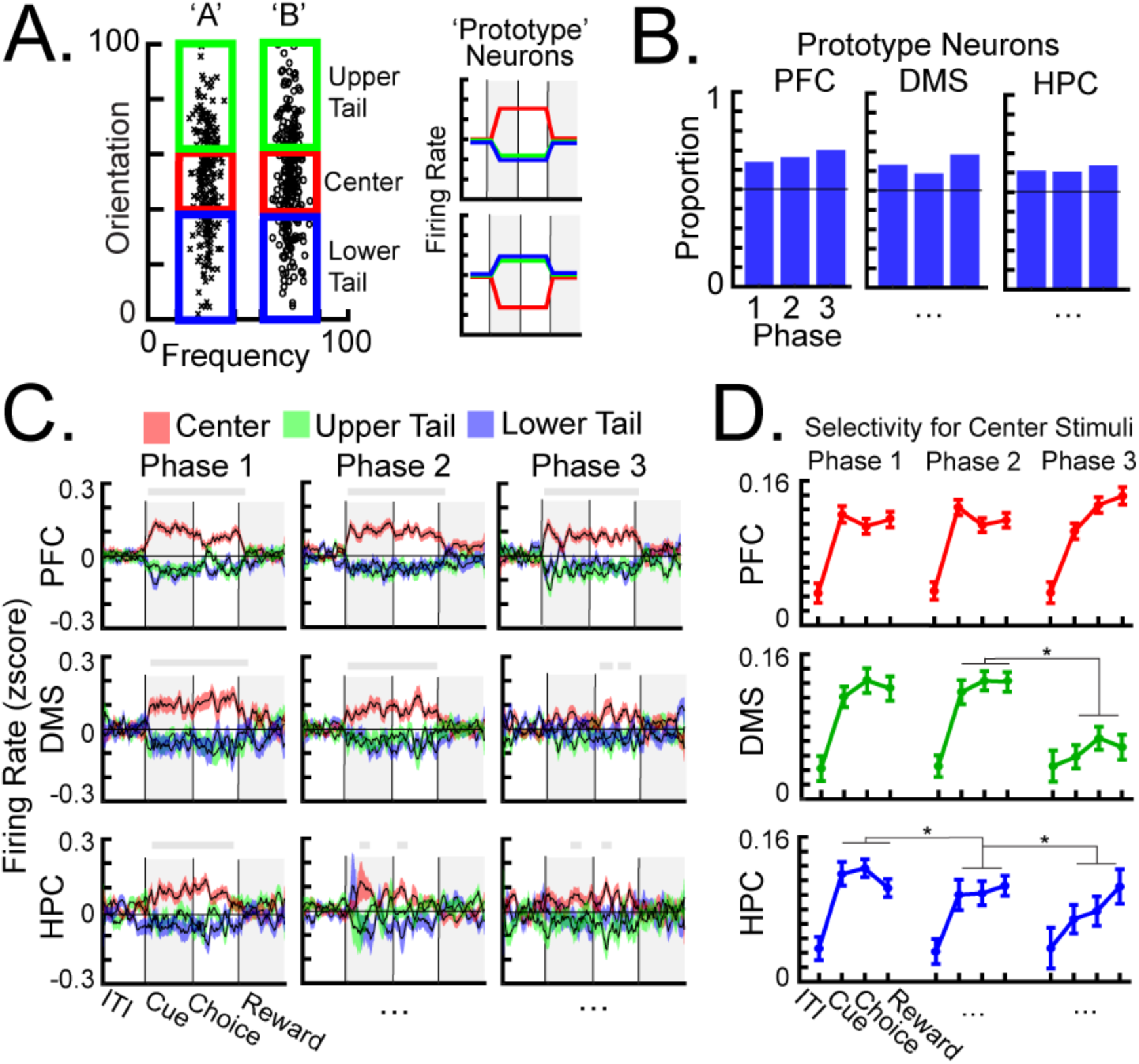
Suppression of irrelevant stimulus information. **A,** Left: The category distributions were divided into three equal sections (i.e., lower tail, center, upper tail). Right: representative “Prototype” neurons had firing rates that peaked (either maximum (top) or minimum (bottom)) for the center stimuli. **B,** The proportion of Prototype neurons in each region and training phase. **C,** Average firing rate of Prototype neurons in each region. The sign of these firing rates was aligned so that all neurons had the same orientation. Gray bars above each plot indicate timepoints in which the firing rates were significantly different between the center stimuli and the tail stimuli. **D,** Selectivity between the center stimuli and the tail stimuli. Selectivity decreased across training phases in the DMS and the HPC, but not the PFC. All error bars indicate *S.E.M*.

### Differentiating stimulus-related and decision-related activity

Our results so far suggest that brain regions (especially the DMS and the HPC) seem to be involved in both categorization and decision-making. Here, we differentiated between these processes and examined how they interacted. First, we used linear regression to correlate each neurons’ firing rates with 1) the category membership of each trial and 2) the rats’ choices (Fig. 4A). This analysis also included incorrect trials, as these trials contained mismatching information about the stimulus on the screen and the rats’ choice. “Category” neurons had firing rates that were more strongly correlated to the category membership of each stimulus, whereas “Decision” neurons had firing rates that were more strongly correlated to the rats’ choices. Generally, there were more Decision neurons than Category neurons (Fig. 4B; PFC: *X*^2^(1, *N* = 1203) = 155.85, *p* < .001; DMS: *X*^2^(1, *N* = 618) = 221.52, *p* < .001; HPC: *X*^2^(1, *N* = 570) = 205.2, *p* < .001). The proportion of Category neurons was larger in the PFC than the other regions (PFC vs. DMS: *X*^2^(1, *N* = 1821) = 28.89, *p* < .001; PFC vs. HPC: *X^2^*(1, *N* = 1773) = 27.55, *p* < .001).

**Figure 4.**
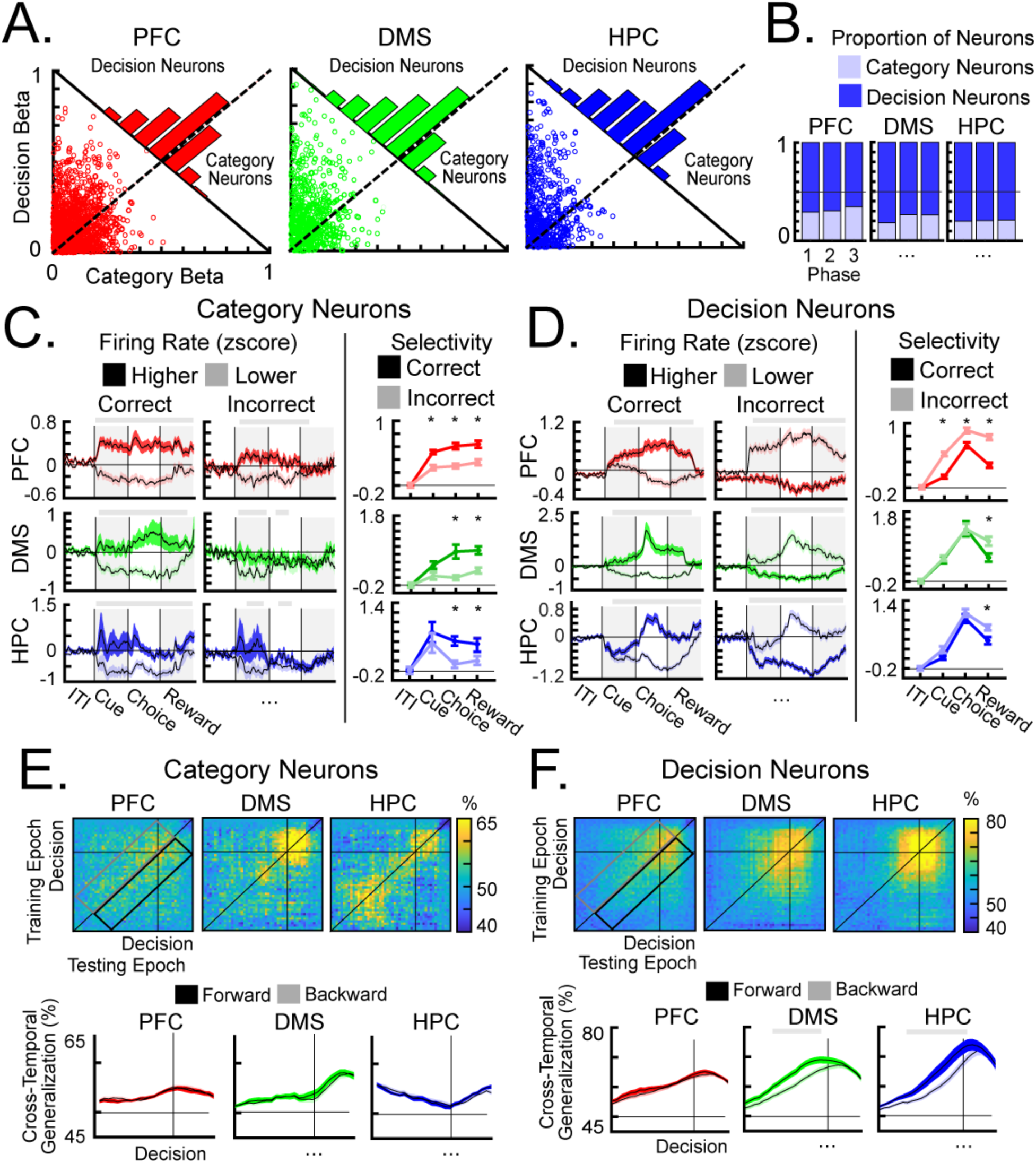
Differentiating stimulus-related and decision-related selectivity. **A,** Correlation coefficients between each neurons’ firing rate and 1) the category membership of each trial (Category beta) and 2) the rats’ choices (Decision beta). “Category” neurons had larger coefficients for the category membership of each trial (below the diagonal). “Decision” neurons had larger coefficients for the rats’ choices (above the diagonal). **B,** The proportion of Category and Decision neurons across training. Most neurons in each region (∼75%) were classified as Decision neurons. There were more Category neurons in the PFC than the other regions. **C-D,** Average firing rates (left) and category-selectivity (right) during correct and incorrect trials of Category neurons (**C**) and Decision neurons (**D**). Gray bars above each plot indicate timepoints in which the firing rates were significantly different between the categories. **C,** Category neurons had weaker category-selectivity during incorrect trials than correct trials. **D,** For Decision neurons, the relative firing rates for the categories were reversed during incorrect trials. **E-F,** Cross-temporal decoding for Category (**E**) and Decision neurons (**F**), aligned to the rats’ decisions. Top: average classifier accuracies for cross-temporal decoding. Points along the diagonal reflect real-time decoding, and points off the diagonal reflect cross-temporal generalization. Regions above the diagonal reflect forward-direction generalization, and regions below the diagonal reflect backward-direction generalization. Bottom: average forward- and backward-direction cross-temporal generalization (the black box and the gray line, respectively) leading up to the rats’ decision. For Decision neurons in the DMS and the HPC, forward-direction generalization was stronger than backward-direction generalization leading up to the rats’ decision. Gray bars above each plot indicate significant differences between directions. All error bars indicate *S.E.M*.

The neural responses of Category neurons and Decision neurons seemed to match their general functions (i.e., stimulus-related vs. decision-related processing). Category neurons had faster selectivity onset than Decision neurons (Supplemental Fig. 6A), suggesting that they were more important for fast-latency categorization. During incorrect trials, Category neurons, and not Decision neurons, had weaker category-selectivity than during correct trials (Figs. 4C&D). This reduction in category-selectivity occurred during the Choice phase and Reward phase in all regions (*p*s < .01) and during the Cue phase in the PFC (*p* = .006). By contrast, Decision neurons in the DMS and the HPC had firing rates that peaked around the rats’ decisions (Supplemental Fig. 6B), suggesting that these neurons were important for decision-making. During incorrect trials, Decision neurons had firing rates that swapped between categories (Fig. 4D). The “Lower” category now elicited a stronger response, and the “Higher” category now elicited a weaker response. This reversal was indicative of the rats’ incorrect choice. Together, these results suggest that Category neurons and Decision neurons had distinct roles during learning.

Decision neurons in the DMS and the HPC seemed to accumulate evidence leading up to the rats’ decisions. We trained Support Vector Machines (SVM) to predict the category membership of each trial using spiking activity from neurons in each region. As expected, category information ramped up and peaked around the rats’ decisions in classifiers trained with Decision neurons, but not Category neurons (Supplemental Fig. 6C). We used cross-temporal decoding to examine how this category information generalized across timepoints (Figs. 4E&F). Cross-temporal generalization was stronger for Decision neurons than Category neurons (Figs. 4E&F), suggesting that category information was maintained across time in Decision neurons. For DMS and HPC Decision neurons, forward-direction generalization (i.e., generalization to future timepoints) was stronger than backward-direction generalization (i.e., generalization to past timepoints) from ∼750 ms before the rats’ decision (Fig. 4F). This asymmetry between forward- and backward-direction generalization is indicative of evidence accumulation and integration processes. For these neurons, category information at a current timepoint became the basis for information during future timepoints. Weaker backward-direction generalization means that additional processing was added after each timepoint. These results suggest that the DMS and the HPC contributed to decision-making by accumulating category information.

### Learning-related changes in theta synchrony

Lastly, we examined the LFPs to assess learning-related changes in inter-regional communication. Category learning was correlated with changes in theta (4-10Hz) oscillations. Average theta power increased across training phases in all regions and all trial events (Supplemental Fig. 7A&B). Peaks in theta power were observed during the Choice phase and the Reward phase (Supplemental Fig. 7A). For the PFC and the HPC, these peaks during the Choice phase were aligned to the rats’ decisions (Supplemental Figs. 7C&D). There was also an alpha/beta (10-25 Hz) component in the HPC during the Reward phase (Supplemental Fig. 8B). Otherwise, there were no meaningful changes in alpha/beta and low gamma frequencies during the trial events and around the rats’ decisions.

Early training sessions relied on DMS-PFC theta Granger causality. We used Granger causality to examine how theta influences flowed between regions (Fig. 5). Throughout training, theta was directional from the DMS to the PFC (Phases 1-3; *p*s < .001). DMS→PFC theta causality was strongest during training Phase 1 and 2 and decreased during Phase 3 (Fig. 5A; Phases 1 and 2 to 3; *p*s < .01). Within each trial, DMS→PFC theta causality was elevated from baseline during the Cue phase and the first half of the Choice phase (Fig. 5A). Causality decreased after the rats’ touch during the Choice phase, without a clear peak (Fig. 5B). These results suggest that DMS→PFC theta interactions were important for stimulus categorization, especially during early learning.

**Figure 5.**
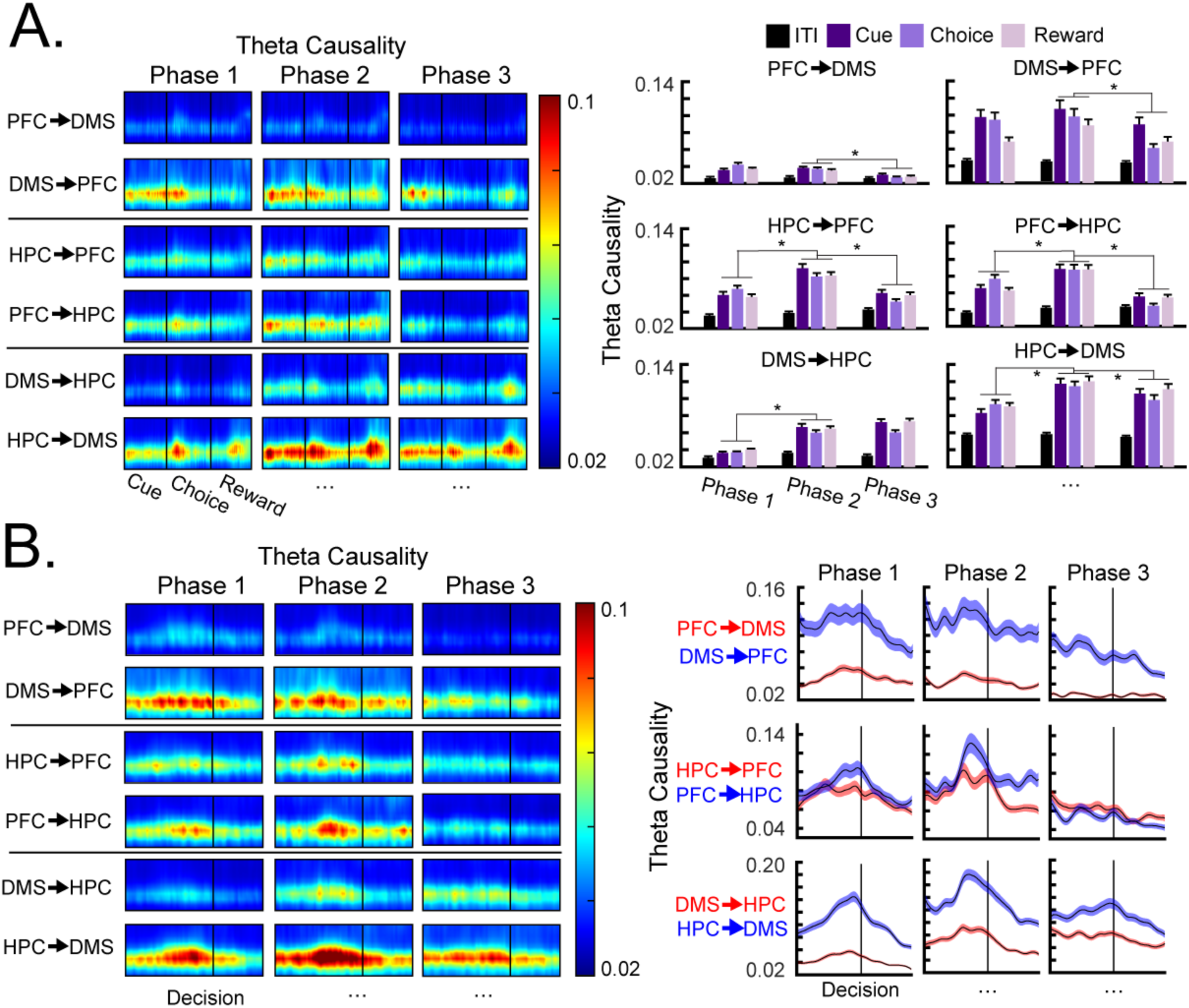
Learning-related changes in theta network communication. **A,** Heatmaps (left) and bar plots (right) of average theta Granger causality between each region pairs. DMS→PFC theta causality was stronger than the opposite direction and decreased across training phases. PFC-HPC theta causality was largely bidirectional, increased from Phase 1 to Phase 2, and decreased from Phase 2 to Phase 3. HPC→DMS theta causality was stronger than the opposite direction, increased from Phase 1 to Phase 2, and decreased from Phase 2 to Phase 3. **B,** Heatmaps (left) and line graphs (right) of average theta Granger causality aligned to the rats’ decisions. DMS→PFC theta causality decreased after the rats’ decision without a clear peak in causality. PFC→HPC theta causality and HPC→DMS theta causality peaked before the rats’ decision during training Phases 1 and 2. All error bars indicate *S.E.M*.

HPC-DMS and HPC-PFC theta causality increased with learning. Throughout training, theta was directional from the HPC to the DMS (Fig. 5A; Phases 1-3; *p*s < .001). Theta influences between the PFC and the HPC were bidirectional (*p*s > .05). HPC→DMS theta causality and bidirectional PFC-HPC theta causality increased from training Phase 1 to Phase 2 and decreased from Phase 2 to Phase 3 (Fig. 5A). Within each trial, PFC-HPC and HPC→DMS theta causality were elevated throughout the trial events, with peaks during the Choice phase and the Reward phase (Fig. 5A). For PFC→HPC causality and HPC→DMS causality, these peaks during the Choice phase were aligned to the rats’ decisions, particularly during training Phases 1 and 2 (Fig. 5B). These results suggest that PFC-HPC and HPC→DMS causality were important during decision-making and increased with learning.

## Discussion

Our results reveal how the PFC-DMS-HPC network learns new visual categories and uses category information to guide decisions in rats. Neuronal category-selectivity was strong throughout training in the PFC and increased with learning in the DMS and the HPC. The DMS and the HPC seemed to be critical for decision-making, with contributions shifting from the DMS to the HPC across learning. In the LFPs, HPC→DMS theta causality and bidirectional PFC-HPC theta causality increased with learning and peaked around the rats’ decisions. By contrast, DMS→PFC was strongest during early training phases and seemed to be more critical for categorization. Together, our results indicate that early learning relied on the PFC and DMS→PFC interactions. After learning, behavior was driven by the HPC, HPC→DMS interactions, and bidirectional PFC-HPC interactions.

### Learning-related changes in neural activity

Early training sessions relied on fast-latency selectivity in the PFC (Figs. 2D&E) and strong DMS→PFC theta causality (Fig. 5). The prefrontal cortex is important for differentiating between category-relevant and irrelevant stimulus information (i.e., attention learning)^5,6,18,23,24^. Attention is closely integrated with categorization and influences how stimuli are encoded and represented^25–27^. We predict that this occurs during early training sessions, so that evolving category representations can emphasize relevant information. Attention learning in the PFC may have been facilitated by DMS→PFC theta interactions. Models of categorization posit that prefrontal-basal ganglia interactions are critical for evaluating category rules^15^. This is supported by neuropsychological experiments in patients with Parkinson’s Disease^28^ and neurophysiological experiments in non-human primates^11^. We hypothesize that the DMS→PFC interactions trained the PFC to differentiate between category-relevant and irrelevant stimulus information.

Representations in the DMS and the HPC became stronger with learning. This was apparent in category-selectivity (Figs. 2D&E) and HPC→DMS theta causality (Fig. 5). Firing rate differences along the irrelevant stimulus dimension decreased across training (Fig. 3), suggesting that these regions learned to filter out the irrelevant dimension. Attention *filtering* may require separate mechanisms than *focusing* attention on relevant information^25,29,30^. Computationally, attention filtering is synonymous with dimensionality reduction^31^ and/or sparse coding^32^, which may be facilitated by the prefrontal cortex^5,33^. Our results suggest that the HPC and DMS contribute to attention filtering during category learning.

The DMS and the HPC were also critical for decision-making. Category-selectivity in these regions (Fig. 2F) and HPC→DMS theta causality (Fig. 5B) peaked around the rats’ decisions. The decision-aligned peaks in category-selectivity increased across training in the HPC and decreased across training in the DMS (Fig. 2F). This implies a shift in contributions from the DMS to the HPC. The striatum is thought to use more “exemplar”-based representations^11,34^, since dopamine neurons can rapidly associate individual stimuli to category labels^35–36^. Conversely, the hippocampus is thought to develop “cluster”-based representations^37–38^. Unlike exemplar representations, clusters form generalizations across category stimuli. These representations are more efficient, because each category can be represented by a small number of clusters that are more easily adaptable to changing category rules or task goals^21,39^. We hypothesize that there was a shift in decision-making from exemplar-based representations in the DMS to cluster-based representations in the HPC.

The shift in contributions from the DMS and the HPC may be mediated by the PFC. After learning, there were increases in PFC-HPC theta causality (Fig. 5). These interactions were largely bidirectional, except during decision-making, in which PFC→HPC theta causality peaked around the rats’ decision (Fig. 5B). These results are consistent with studies showing increased prefrontal-hippocampal synchrony memory inference tasks^40–41^. Models of categorization posit that cluster-based representations in the hippocampus are built via prefrontal-hippocampal interactions^16,17^. For example, the prefrontal cortex decides how new stimuli are integrated with existing representations and biases representations according to current task goals. In our task, we hypothesize that the PFC influences categorization and decision-making in the HPC. This occurs after the PFC has differentiated between relevant and irrelevant stimulus information.

Finally, category-selectivity in the HPC as well as PFC-HPC and HPC→DMS theta causality decreased from training Phase 2 to Phase 3 (Fig. 5). This may reflect a shift in contributions from networks involved during category learning vs. retention. For example, categorization behavior in fully trained categories becomes independent of the regions involved during learning^42–43^. Further experiments can examine how brain networks shift as the behavior becomes more automatic.

### Differentiating between stimulus-related and decision-related activity

We used linear regression to differentiate between Category neurons that tracked the category membership of each trial and Decision neurons that tracked the rats’ choices (Fig. 4). Category and Decision neurons were present in all regions, suggesting that each region contained a mixture of stimulus-related activity and decision-related activity. This also rules out the alternative hypothesis that the category-selectivity observed in Figure 2 was driven solely by premotor activity. During error trials, Category neurons had reduced category-selectivity, indicative of weaker encoding of the current stimulus. By contrast, Decision neurons had firing rates that swapped between categories, indicative of the rats’ incorrect choice. Buetfering et al., (2022)^44^ found similar results in the mouse primary somatosensory cortex: some neurons correlated with the identity of the presented stimulus, whereas other neurons correlated with the mice’s decisions.

Cross-temporal decoding revealed that Decision neurons in the DMS and the HPC were important for decision-making by accumulating category information before the rats’ decisions (Fig. 4F). Evidence accumulation has been demonstrated in multiple brain regions, including the dorsal striatum^45^ and the hippocampus^46^. Nieh et al., (2021)^46^ found that the hippocampus may integrate accumulate evidence with spatial information. In our task, the HPC (and HPC→DMS) may accumulate category information and transform that information into a behavioral response (i.e., a touch to the left or right report key). Future experiments can more directly test these functions using tasks specifically designed to probe evidence accumulation^47^, and/or designs in which categorization and decision-making are separated in time (i.e., a delayed-match-to-category design^4^).

It should be noted that although we split the neurons into Category and Decision groups, the correlation coefficients were largely continuous from “purely stimulus-related” to “purely decision-related”. This suggests that Category neurons and Decision were not distinct subpopulations of neurons, but rather two extremes along a continuum. This is consistent with the concept of “mixed selectivity,” in which each neuron can contribute to a combination of task features^48–49^. Mixed selectivity is an important feature of cognition that creates high-dimensional representations that are flexible and robust to noise.

### The role of inhibition

The majority of DMS and HPC neurons showed decreased activity during the Cue phase and Choice phase (Supplemental Fig. 3). This seemed to be functional, as the proportion of inhibitory HPC neurons increased with learning. Inhibition in this task may be beneficial for suppressing irrelevant category representations and/or actions before the current stimulus is fully encoded^50^. This could be mediated via local inhibitory circuits^51–53^ and/or top-down control from the PFC^54–55^. For example, Malik et al., (2022)^56^ demonstrated that the PFC exerts top-down control through long-range GABA projections that affect inhibitory circuits in the HPC.

## Conclusions

Together, our results provide a framework that describes how the PFC-DMS-HPC network interacts when learning new categories. These results provide valuable insight for models of categorization and are informative to understand the mechanisms underlying neurological disorders that affect category learning, including patients with Parkinson’s Disease^10^, autism^57^, dementia^58^, and others. These results also shed light on how rats generalize across stimuli to learn complex task structures. This is relevant for the development of cognitive maps^59–60^ and schemas^61^.

## Materials and Methods

### Touchscreen Apparatus

All training sessions were contained within an experimental chamber outfitted with a touchscreen (36 × 41 × 36 cm). The right wall of the chamber contained a computer monitor (Model 1550V, NEC, Melville, NY) that presented visual stimuli to the rats. A touchscreen (15-in, Elo Touch Systems, Fremont, CA) overlaid the monitor to allow the rats to interact with the visual stimuli. The opposite chamber wall contained a food tray (6.5 × 13 × 4.5 cm). Food pellets were delivered to the rats via a rotary pellet dispenser (Med Associates Inc., Georgia, VT, model ENV-203IR) controlled by an electrical board (Model RS-232, National Control Devices, Osceola, MO). Custom-written MATLAB scripts (MathWorks, Natick, MA) controlled all experimental procedures. A house light was positioned above the food tray and was always on during sessions. A small hole was drilled into the ceiling of the chamber to allow access to the tether. White noise in the room was used to minimize distractions. Finally, the rats’ behavior was monitored by a camera (model ELP-USB100W05MT-RL36) mounted to the ceiling.

### Pre-Training Procedures

All procedures were approved by the Institutional Animal Care and Use Committee at the University of Iowa. Rats arrived at the animal colony and were given an acclimation period with unlimited access to food and water. After one week, food was restricted to increase motivation. Weights were recorded daily to ensure their weight did not go below 85% of their free feeding weight. Rats were handled daily for one week to reduce the stress of interacting with experimenters. Then, rats underwent a “cart training” procedure to further habituate to the lab environment. During this procedure, rats were encouraged to forage for food pellets on top of an open field (i.e., a laboratory cart). Finally, rats completed a shaping procedure within the experimental chambers to learn to interact with the touchscreens. Each phase of the shaping procedure became increasingly similar to the trial procedure used during training sessions, without exposing the rats to the category stimuli (see Broschard et al., (2020)^62^ for more details).

### Categorization Task

The category stimuli in this experiment were Gabor patches containing black and white gratings. Across stimuli, these gratings varied in their spatial frequency (0.2532 cycles per visual degree to 1.2232 cpd) and orientation (0 radians to 1.75 radians). The ranges of these dimensions are within the perceptual limits of Long-Evans rats in touchscreen-based tasks^63^ and have been used in multiple rat category learning experiments^18,19,20,64^. A two-dimensional stimulus space was created from these dimensions so that category stimuli could be sampled and generated (Fig. 1A). Each dimension was linearly transformed so that the axes of the stimulus space had a common range (i.e., 0 to 100).

Categories were created by positioning normal distributions onto this two-dimensional stimulus space (Fig. 1A). The points within one distribution (*µ_X_* = 30, *σ_X_* = 2.5, *µ_Y_* = 50, *σ_Y_* = 20) constituted the members of category ‘A’, and the points within the other distribution (*µ_X_* = 70) constituted the members of category ‘B’. These distributions were perpendicular to the spatial frequency dimension and parallel to the orientation dimension. Because of this alignment, the categories were typically learned by attending to the spatial frequency dimension and ignoring the orientation dimension. The distributions were regenerated before each session; therefore, the rats were shown novel stimuli on every training trial. This ensured the rats could not learn the task by simply memorizing a few exemplars.

For simplicity, we did not include a counterbalancing task in which orientation was the relevant dimension and spatial frequency was the irrelevant dimension. Across many experiments, we have found that rats learn both tasks at the same rate and to equivalent levels^18,18,20,62^.

### Trial Procedures

Each training session contained 80 trials. Each trial was self-initiated by touching the ‘Star’ cue positioned at the center of the screen (Start phase; Fig. 1B). Then, a unique category stimulus was sampled from the category distributions and was presented at the center of the screen for two seconds (Cue phase). The rats were required to touch this stimulus at least once. This ensured attention was directed towards the screen. Next, report keys appeared on the left and right sides of the screen for two seconds (Choice phase). The rats touched either report key, depending on the category membership of the stimulus presented during the Cue phase. The rats learned a spatial mapping of the categories. For example, the left report key was correct for members of category ‘A’, and the right report key was correct for members of category ‘B’. This was counterbalanced across rats. A food pellet was delivered after correct responses (Reward phase). After each incorrect response, the trial was aborted and repeated without reinforcement. Inter-trial intervals ranged from five to ten seconds. Trials were aborted if the rats did not complete their touch requirements (i.e., one touch during Cue phase and one touch during the Choice phase) within the allotted time constraints. This rarely occurred.

The report keys were direct copies of the category stimulus presented during the Cue phase. This was not necessary, since the categories were mapped to spatial locations and the identity of each report key was irrelevant. However, we have found that this improves the rats’ performance. We suspect this procedure improves S-R learning by minimizing the working memory delay between the presentation of the stimulus and the rats’ decision.

### Training Phases

Rats were given fifteen training sessions to learn the category task. This was chosen based on previous experiments that have found that rats typically reach 75% accuracy within ten training sessions^18,19,20,64^. We divided the training sessions into three training phases (i.e., Phase 1, Phase 2, and Phase 3) to examine neural activity at different timepoints in learning. Each phase averaged across five training sessions. To ensure that the selection of these training phases did not induce unnecessary biases in the results, we reorganized the data according to session accuracy (Supplemental Fig. 8; i.e., <60%, 60-70%, and >70%). Importantly, learning-related changes in firing rate and LFP connectivity were qualitatively similar between methods.

### Surgery

Each rat underwent stereotaxic surgery under isoflourane (1% - 4%) anesthesia to implant a custom-built, 3D-printed microdrive. Prior to implant, each tetrode’s impedance was lowered to 150-300 kΩ by electroplating it in a gold non-cyanide solution (NanoZ, Neuralynx). Each microdrive was aligned so that exit tips were positioned over the PFC (AP: 3.0; ML: 0.7 DV: -3.5), DMS (AP: -0.4; ML: 2.25; DV: -4.5), and HPC (AP: -3.8; ML: 2.5; DV: -3.2). Each of the eight recording tetrodes moved independently and each recording site had its own reference. After removing dura mater, tetrodes were lowered 1.0 mm into the brain. After surgery, rats were placed on a heating pad until awake and mobile to prevent hypothermia. Meloxicam (1 mg/ml) was administered as an analgesic during surgery and 24 hours after surgery to improve recovery. Rats were allowed at least one week to fully recover.

### Electrophysiological Recordings

After recovery, tetrodes were slowly lowered in 0.25 mm increments. Recording tetrodes were lowered to their target site (PFC: -3.5 mm; HPC: -3.2 mm; DMS: -4.5 mm). Once within the target region, small adjustments were made (e.g., 0.1 mm) until single units were isolated and stable on the majority of the recording tetrodes. The reference tetrodes were lowered to about 1.0 mm above the target loci. Signals from each region were referenced by a unique reference tetrode to minimize volume conduction. Data were amplified and digitized using data acquisition software (Cheetah, Neuralynx). LFPs were recorded at 30 kHz and down sampled to 1 kHz. Spiking activity was recorded at 32 kHz.

### Histology

After all behavioral testing, the final position of each recording tetrode was marked by electrolytic lesions (10 µA current for 10 s). Then, each rat was given a lethal dose of euthanasia solution (sodium pentobarbital) and was perfused with ∼150 mL PBS and ∼150 mL formalin. Brains were stored at 4 degrees C. A sliding microtome made 50 µm coronal sections of the target brain regions. Slides were stained with thionin and cover slipped. Once dry, each slide was observed under a light microscope to examine the location and spread of each marker lesion. The boundaries of the PFC, HPC, and DMS were defined according to Paxinos & Watson, (1998)^65^.

### Statistical Analysis

Linear mixed effects modeling (R, version 3.4.2) was used to assess learning-related changes in behavior. Models included fixed effects for training phase and a quadratic function across data points. Models also included random effects for slope, intercept, and the quadratic function. A model simplification strategy was used to find the simplest model that fit the model^64^. We started with the full model and systematically removed random effects one at a time. This continued until the estimates were significantly different from the larger model before it. Chi square tests were used to assess whether proportions were significantly different between groups.

Non-parametric statistics were used for all neural measures, as we could not assume these data were normally distributed. Each neuron or LFP was treated as an independent observation. Randomizations were performed across all samples and were repeated 10,000 times to obtain a *p* value.

### Spiking Analyses

#### Preprocessing

Spikes from single units were isolated off-line using the cluster cutting software (MClust 4.4). Multiple parameters, including peak, width, height, and energy associated with the waveforms were used to isolate single units. Inter-spike interval histograms confirmed that no single unit spikes occurred within a 2 ms window of each other. Neurons with a grand average firing rate less than 1.0 Hz were removed. Unless otherwise noted, analyses only included correct trials.

#### Firing Rate

Firing rates were calculated by summing binary spiking activity into 50-ms nonoverlapping bins. Firing rates were normalized by converting these values to z-scores relative to baseline (i.e., two seconds of data during the ITI period). Firing rates were separated for category ‘A’ trials and category ‘B’ trials. For each neuron, we organized the firing rates depending on which category had larger firing rates during the Cue phase and the Choice (i.e., “Higher” and “Lower”). Organizing the categories in this way maximized the firing rate differences between categories.

#### Category-Selectivity

Category-selectivity was quantified using the category tuning index (CTI) described by Freedman et al., (2001)^4^. This index compared firing rates for within-category trial pairs (e.g., two category ‘A’ trials) and between-category trial pairs (e.g., a category ‘A’ trial vs. a category ‘A’ trial). Specifically,

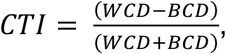

where WCD was the average difference in firing rate for all within-category trial pairs, and BCD was the average difference in firing rate for all between-category trial pairs. CTI values ranged from 0 to 1, where 0 indicated no category selectivity, and 1 indicated perfect selectivity. CTI values were calculated during each trial event (i.e., two seconds of data). For analyses that examined continuous changes in CTI, we used a moving window of 200 ms with 100 ms overlap between each bin.

We found that the CTI values were inflated when fewer trials were included in the calculation. This was an issue for analyses that compared selectivity for conditions with uneven numbers of trials (e.g., correct vs. incorrect). For those analyses, we estimated category-selectivity by subtracting the mean firing rate between trials at each category at each timepoint.

#### Selectivity Onset

CTI onset measured the latency of selectivity onset. For each neuron, CTI values were calculated iteratively, averaging across larger timepoints of the trial on each iteration. Specifically, the first iteration used data from the first 100 ms of the Cue phase, the second iteration used the first 150 ms of the Cue phase, and so on. Iterations continued (adding 50 ms of data) until the end of the Choice phase was reached. This resulted in a curve of CTI values that generally increased monotonically until reaching a maximum value. We normalized this curve from 0 to 1, representing the neuron’s smallest and largest level of category-selectivity. Selectivity onset was defined as the latency to reach at least 50% of the maximum value. Similar results were obtained when other thresholds were used (i.e., 25% and 75%).

#### Selectivity Along the Irrelevant Stimulus Dimension

To compare neural responses along the irrelevant stimulus dimension, we subtracted off the mean firing rate of each category at every timepoint within the trial. This removed between-category variances (i.e., category-selectivity) and isolated variances within each category.

We divided the category distributions into three divisions along the irrelevant dimension. “Center” stimuli were defined as stimuli within +/- 0.5 SD of the category mean along the irrelevant dimension (i.e., orientation values between 40 and 60). “Upper Tail” stimuli were greater than +0.5 SD of the category mean (i.e., orientation values larger than 60), and “Lower Tail” stimuli were lower than -0.5 SD of the category mean (i.e., orientation values lower than 40). These divisions were chosen so that each trial type had roughly equal trial numbers.

“Prototype” neurons had firing rates (averaged across the Cue phase and the Choice) that peaked (either maximum or minimum) for the Center stimuli and were smaller (or larger) for stimuli at both tails. We reversed the sign so that all Prototype neurons had firing rates that peaked (i.e., maximum) for the Center stimuli. This allowed us to average activity across neurons.

#### Correlating firing rates with task features

We used linear regression to correlate each neurons’ firing rate with 1) the category membership of each trial (i.e., category ‘A’ vs. category ‘B’) and 2) the rats’ decision (i.e., a leftward response vs. a rightward response). It was critical to include incorrect trials in this analysis, since these trials contained mismatching category and decision information. We assumed that each neuron’s firing rate activity was a weighted sum of the category and decision task variables. Specifically,

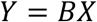

where *Y* was the neuron’s firing rate of each trial averaged across the Cue phase and the Choice phase, *X* was a matrix with size *trials* x *task features* and contained trial-by-trial binary information about the category membership of each trial and the rats’ decision, and *B* was a [1 x 2] vector with correlation coefficients for the category and decision terms. For each region, we estimated vector *B* by using Ordinary Least Squares. Specifically,

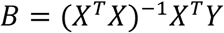

We computed the absolute value of each correlation coefficient, since we were interested in the strength of the correlation rather than its direction. “Category” neurons were defined as neurons with a larger correlation coefficient for the category membership of each trial than the rats’ decision. “Decision” neurons were defined as neurons with a larger correlation coefficient for the rats’ decision than the trials’ category membership.

#### Cross-Temporal Decoding

We trained Support Vector Machines^66^ (SVM) to predict the category membership of each trial using firing rate activity binned at 100 ms. To ensure this analysis had a sufficient number of neurons, we combined neurons recorded from all sessions within a given training phase (and within the same rat). The classifiers were trained using the MATLAB function *fitcsvm* with the *RBF* kernel function. We tested the classifiers on 20% of trials that were not included in the training set. We minimized bias in the classifiers by randomizing the trials used for the training and testing sets on each iteration and averaging the classifier accuracies across 100 iterations. We ensured both the training and testing sets contained an equal number of category ‘A’ and category ‘B’ trials.

Cross-temporal decoding extended this analysis by testing the classifiers on data from all timepoints within the trial^67^. This procedure created a matrix of classifier accuracies, where the trial timepoint used to train the classifier was on one axis and the trial timepoint used to test the classifier was on the other axis. Real-time decoding appeared along the diagonal, and cross-temporal generalization appeared off the diagonal.

### LFP Analyses

#### Preprocessing

Outlier tetrodes were identified by computing the grand mean raw signal for each tetrode. Tetrodes at least four standard deviations above each region’s population mean were removed. This rarely occurred, but ensured the effects were not biased by outliers. We removed evoked potentials from each tetrode by subtracting off the trial-averaged raw signal. Only correct trials were used for each analysis.

#### Power

Power information was calculated by bandpass filtering (Butterworth) the raw LFP signals at each frequency (i.e., 4-40 Hz) and computing the Hilbert transformation on the bandpass filtered data. Power at each timepoint was computed by squaring the absolute value of this complex signal. Scores were normalized by converting raw values to z-scores relative to baseline (i.e., three seconds of data during the ITI period).

#### Granger Causality Analysis

Granger causality analysis^68^ was used to estimate the directionality of theta oscillations between tetrode pairs. Briefly, Granger causality predicts future values of a timeseries using past values. Timeseries *X* ‘Granger-causes’ timeseries *Y* if adding past values of *X* increases the prediction of future values of *Y.* For this analysis, we used the *FieldTrip* toolbox in MATLAB. All comparisons used a model order of 15, similar to Fiebelkorn et al., (2018)^69^. Qualitatively similar results were found using model orders of 13 and 17. The data were down sampled to 250 Hz to reduce the computation time. The data were detrended to minimum effects caused by nonstationary drifts in the data^70^.

## Aligning trials to the rats’ decision

We examined activity related to decision-making by aligned trials to the rats’ touch during the Choice phase. Each analysis was calculated across four seconds of data (i.e., three seconds before the Choice touch and one second after the Choice touch). We excluded trials in which the Choice touch occurred within one second of the onset of the Reward phase. This ensured that the analysis did not include feedback-related activity.

## Acknowledgements

National Institutes of Health Grant P01-HD080679 to J.H.F. National Research Foundation of Korea NRF-2024S1A5A8019683 to J.K.

## Data availability

All data will be made available upon acceptance.

## Author Contributions

*MBB*: Conceptualization, Data curation, Investigation, Formal analysis, Software, Methodology, Visualization, Validation, Writing-original draft, Writing-review & editing

*JK*: Conceptualization, Data curation, Investigation, Formal analysis, Software, Methodology, Writing-original draft, Writing-review & editing

*HEH*: Resources, Methodology, Investigation, Writing-review & editing

*SJF*: Resources, Methodology, Investigation, Writing-review & editing

*JHF*: Conceptualization, Investigation, Funding acquisition, Project administration, Supervision, Writing-original draft, Writing-review & editing

---

**Supplemental Figure 1.**
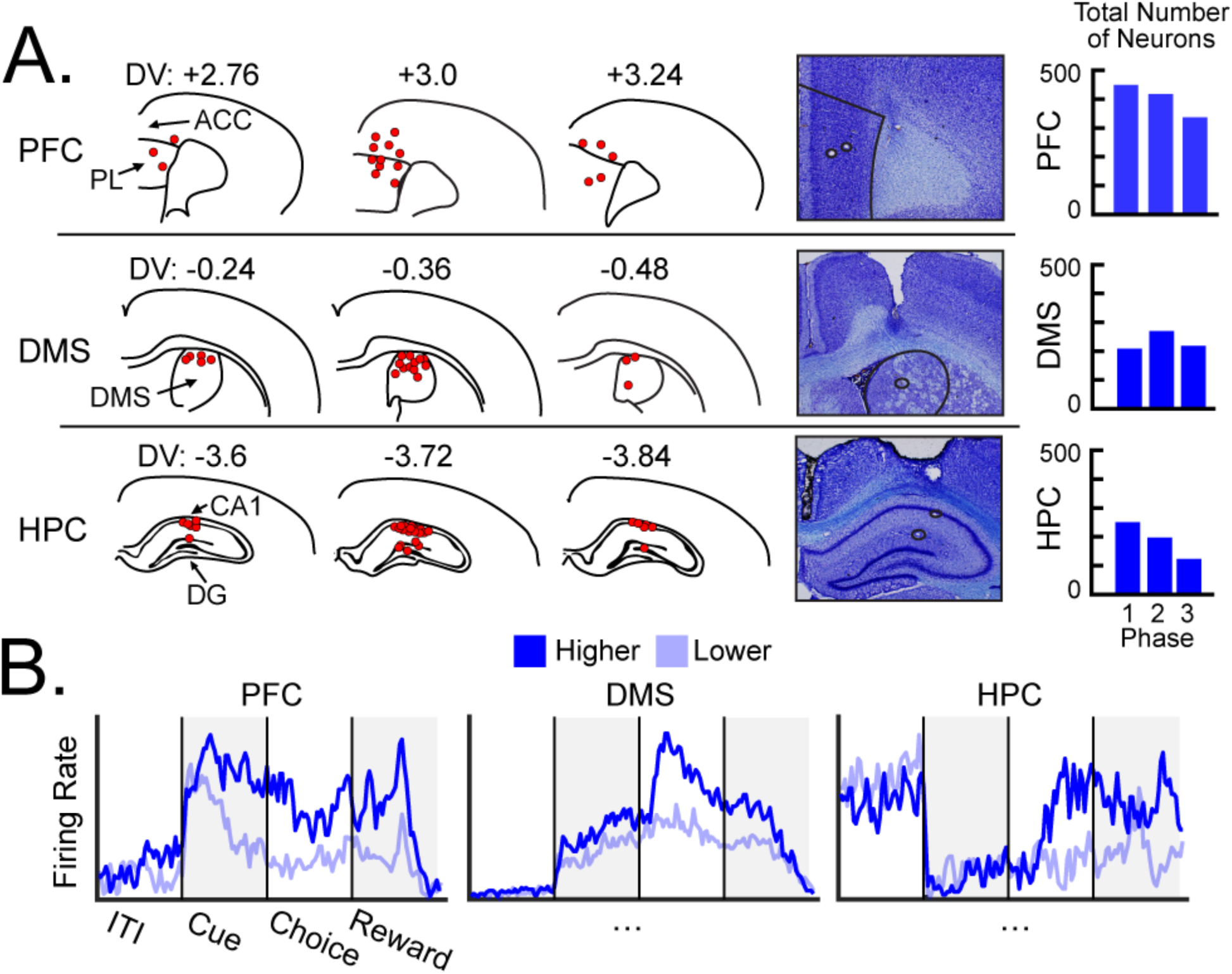
Recording locations. **A,** Left: the placement of recording tetrodes was determined via electrolytic lesions. In the PFC, tetrodes spanned dorsal portions of the rostral anterior cingulate (ACC) and ventral portions of the prelimbic cortex (PL). In the DMS, tetrodes were concentrated in the dorsal portions of the DMS. In the HPC, tetrodes were concentrated in the CA1 and extended into the dentate gyrus (DG). Middle: example tetrode placement. Right: the number of neurons in each region and training phase. **B,** Firing rates of representative neurons recorded during a single session, separated by the neurons’ ‘Higher’ and ‘Lower’ categories.

**Supplemental Figure 2.**
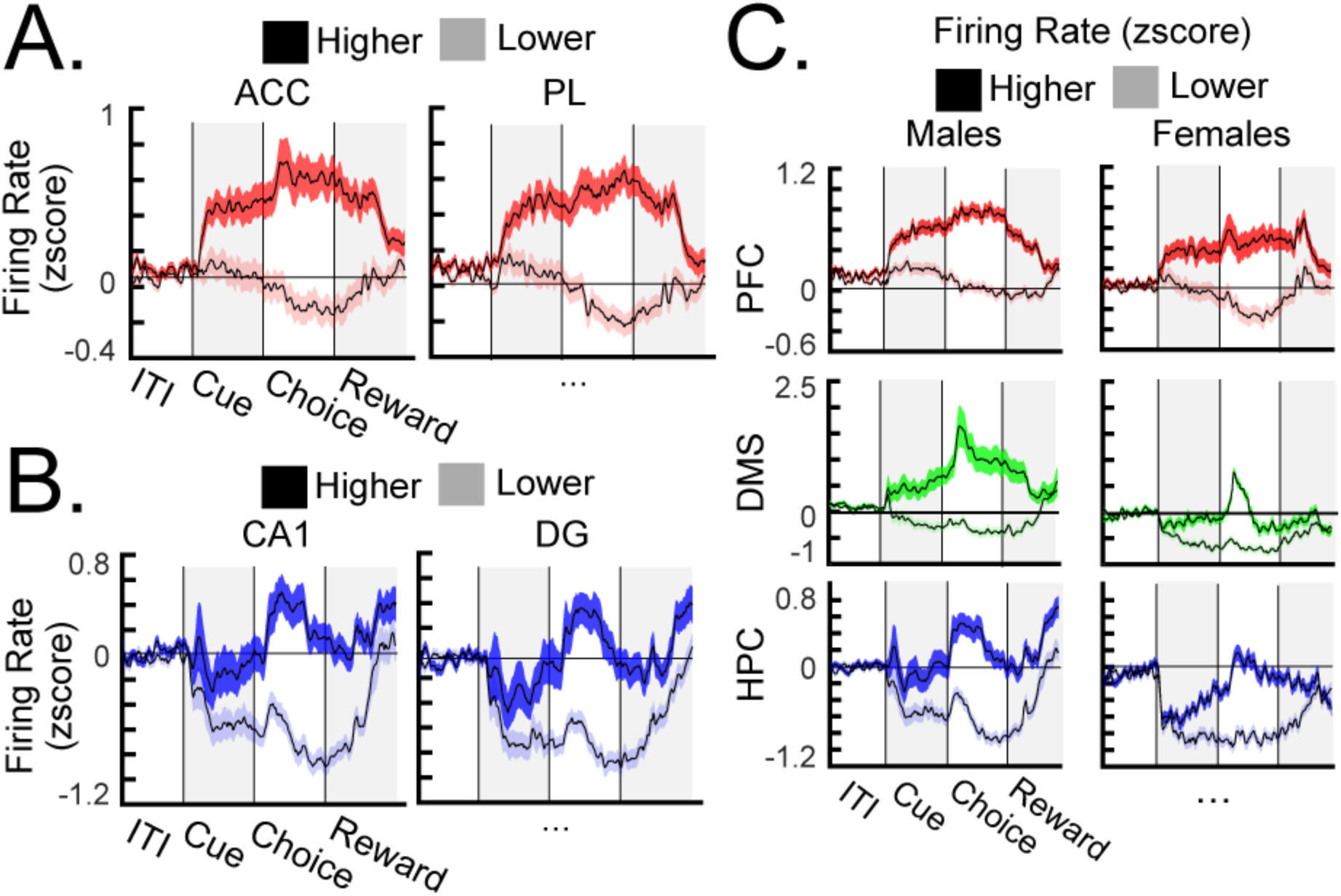
Firing rates. **A,** Average firing rates of PFC neurons centered in the ACC (left) and the PL (right). **B,** Average firing rates of HPC neurons centered in the CA1 (left) and the DG (right). **C,** Average firing rates of neurons for male rats (left) and female rats (right). All error bars indicate *S.E.M*.

**Supplemental Figure 3.**
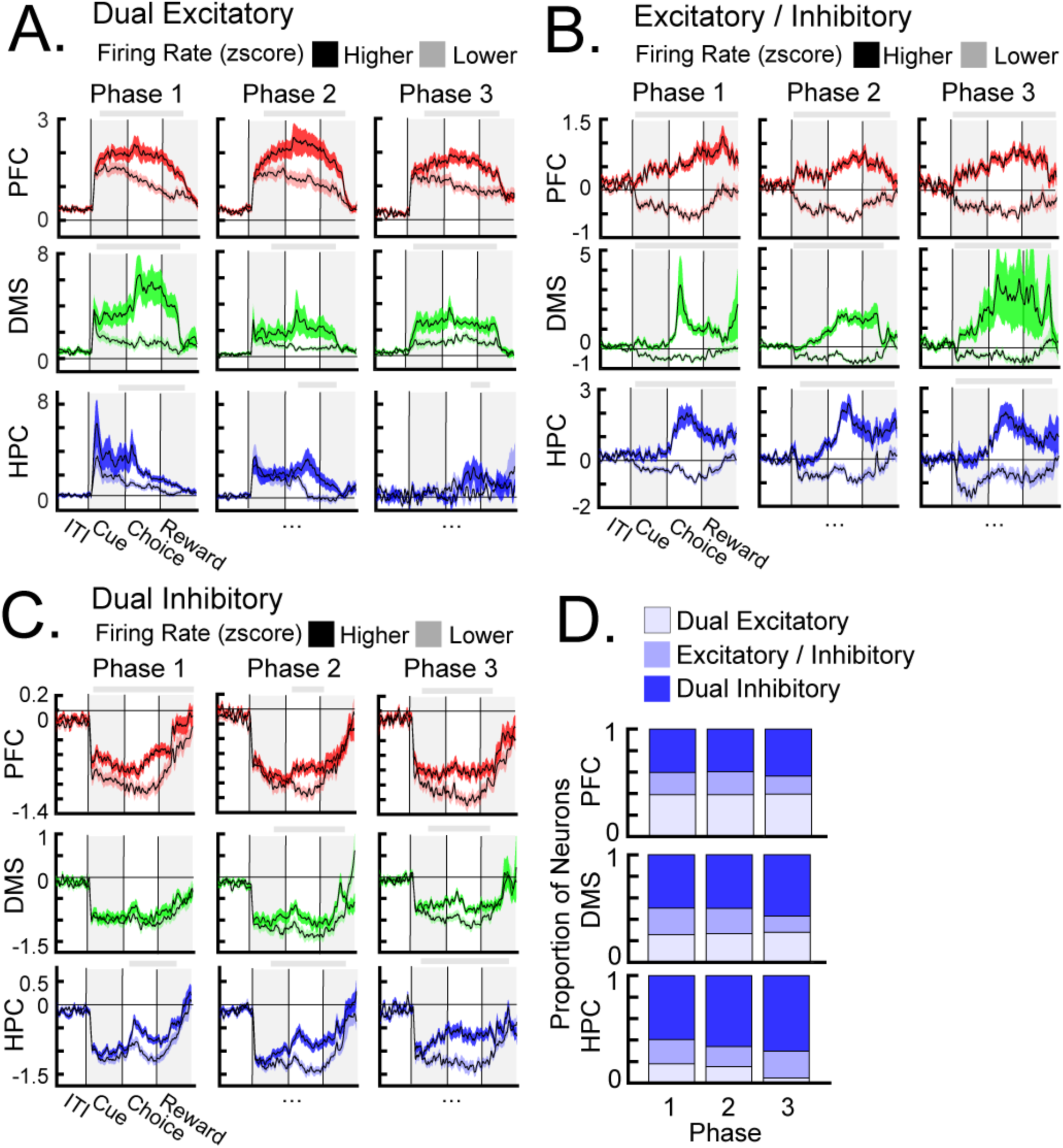
Neurons were separated into three subdivisions according to their firing rate patterns. “Dual Excitatory” neurons had above-baseline firing rates for both categories. “Excitatory / Inhibitory” neurons had above-baseline firing rate for one category and below-baseline firing rate for the other category. “Dual Inhibitory” neurons had below-baseline firing rates for both categories. **A-C,** Average firing rates of Dual Excitatory neurons (**A**), Excitatory / Inhibitory neurons (**B**), and Dual Inhibitory neurons (**C**), separated by each neurons’ ‘Higher’ and ‘Lower’ categories. **D,** The proportion of neuron types across training phases. The proportion of Dual Inhibitory neurons was larger than the proportion of Dual Excitatory neurons in the DMS and the HPC. All error bars indicate *S.E.M.* Gray bars above each plot indicate timepoints in which firing rates were significantly different between categories.

**Supplemental Figure 4.**
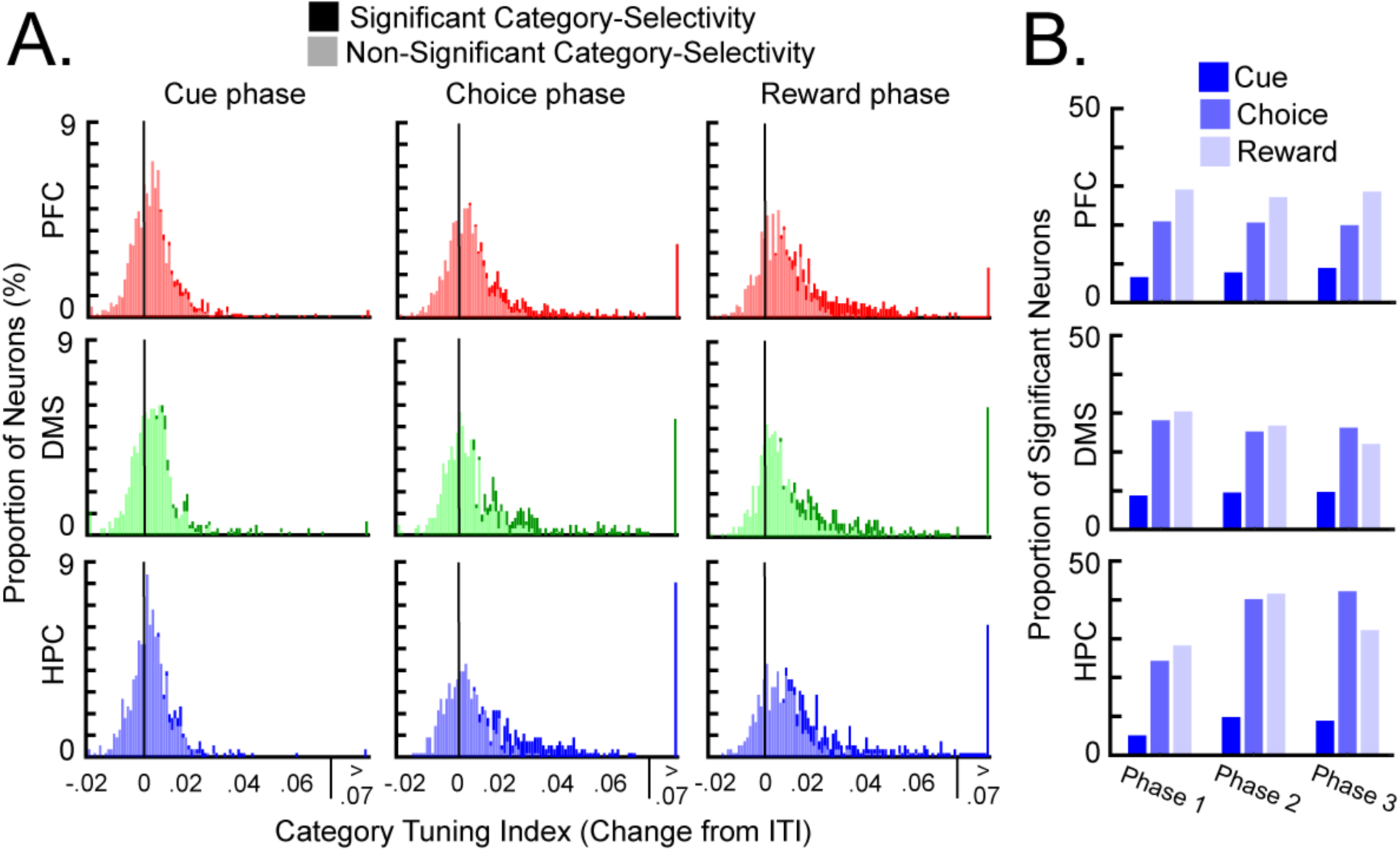
A small proportion of neurons showed significant changes in category-selectivity from baseline. **A,** The distribution of CTI values (subtracted from baseline) of each neuron at each trial event. Neurons with CTI values that that were significantly larger from baseline (*p* < .010) were largely positioned at the right tail of the distributions (darker color). **B,** The proportion of neurons in each region, training phase, and trial event with CTI values that were significantly larger from baseline.

**Supplemental Figure 5.**
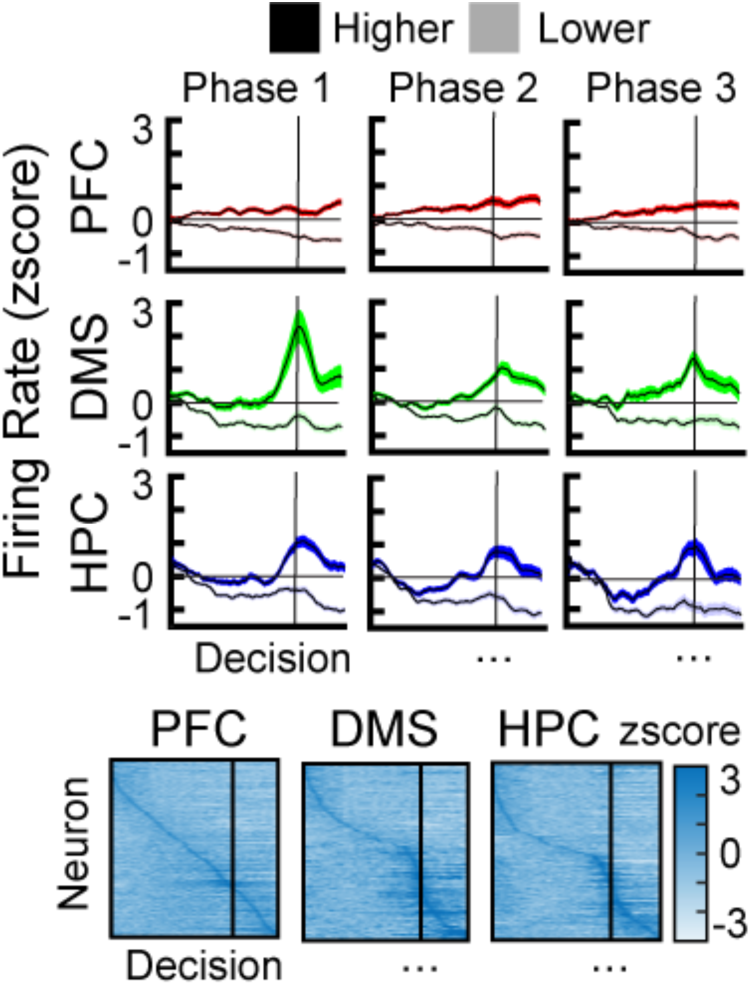
Firing rates aligned to the rats’ choices. Top: the average firing rate of DMS and HPC neurons peaked around the rats’ choices. Bottom: a large proportion of DMS and HPC neurons had firing rates that peaked around the rats’ choices. Error bars indicate *S.E.M*.

**Supplemental Figure 6.**
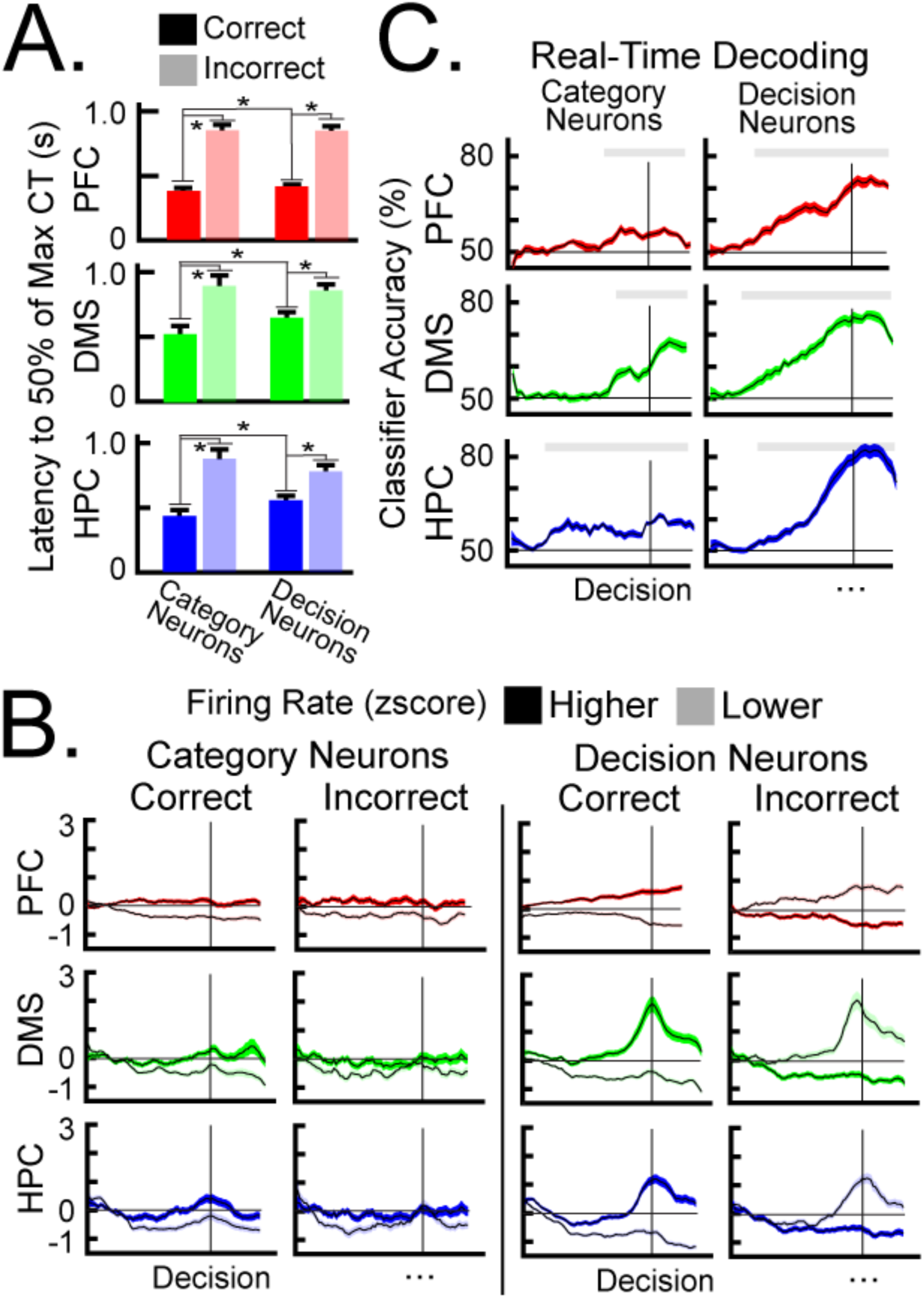
Differentiating stimulus-related and decision-related selectivity. **A,** Selectivity onset measured the latency for each neuron to reach 50% of its maximum category-selectivity. Selectivity onset was faster for Category neurons than Decision neurons. For both Category and Decision neurons, selectivity onset was faster during correct trials than incorrect trials. **B,** Average firing rate of Category neurons (left) and Decision neurons (right) aligned to the rats’ decisions. Firing rates of Decision neurons, and not Category neurons, peaked around the rats’ decisions. For Decision neurons, and not Category neurons, the relative firing rates of the categories reversed during incorrect trials. **C,** SVM classifiers were trained to predict the category membership of each trial. Classifier accuracies for Decision neurons (right) but not Category neurons (left) ramped up before the rats’ decisions and peaked around the decision. All error bars indicate *S.E.M*.

**Supplemental Figure 7.**
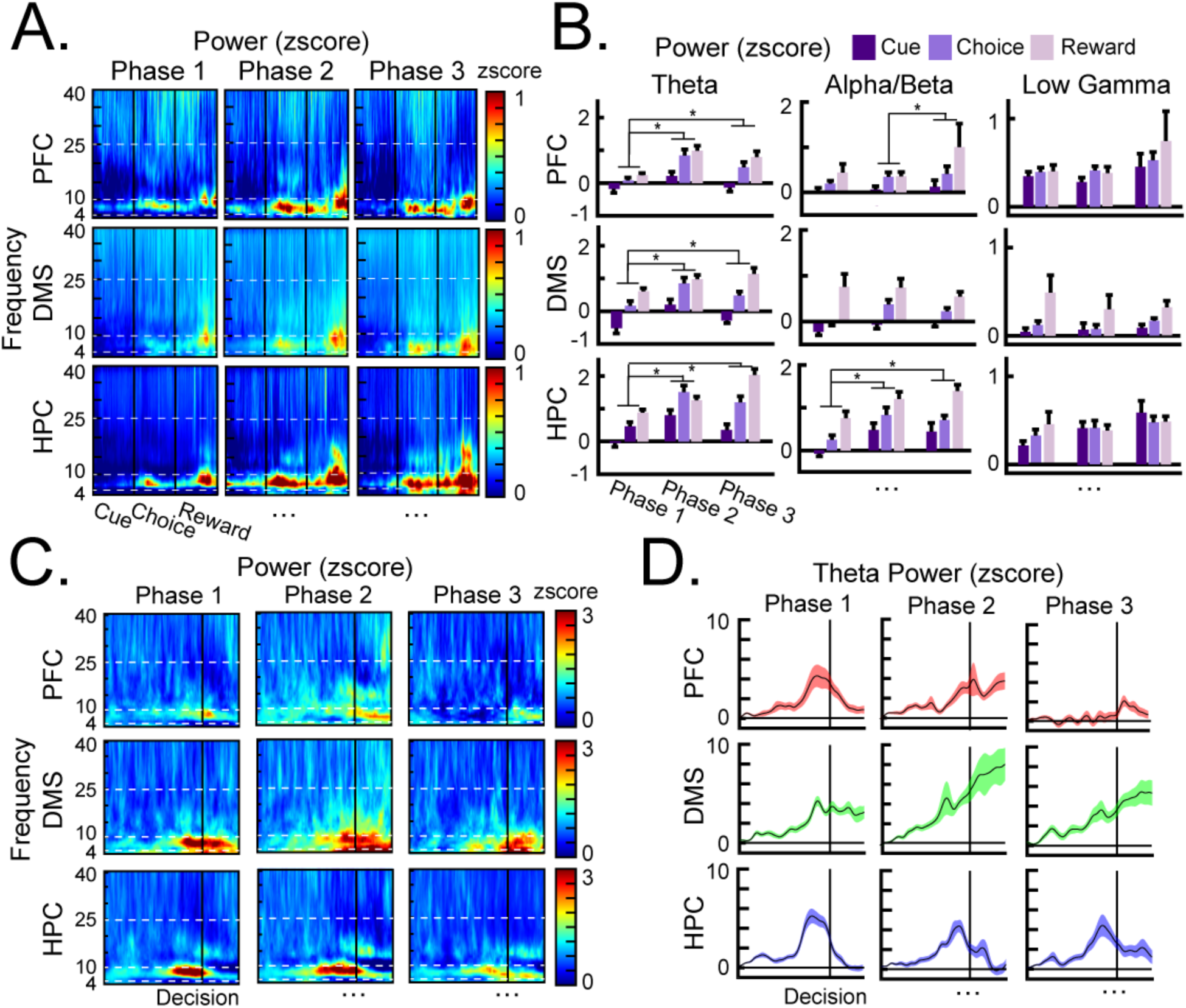
Category learning was governed by theta oscillations. **A,** Heatmaps of averaged power. White dotted lines separated theta (4-10Hz), alpha/beta (10-25Hz), and low gamma (25-40Hz) frequency bands. Peaks in theta power were observed in each region during the Choice phase and Reward phase. **B,** Average power for each frequency band. Theta power increased across training phases in each region. Alpha/beta power increased across training phases in the HPC. There were no meaningful changes in low gamma power. **C,** Heatmaps of average power aligned to the rats’ decision. White dotted lines separated the theta, alpha/beta, and low gamma frequency bands. **D,** Average theta power around the rats’ decision. PFC theta power peaked before the rats’ decision during training Phase 1. DMS theta power peaked before the rats’ decision during training Phase 1. For training Phases 2 and 3, DMS theta power increased through the trial events, without a clear peak. HPC theta power peaked before the rats’ decision during all training phases. All error bars indicate *S.E.M*.

**Supplemental Figure 8.**
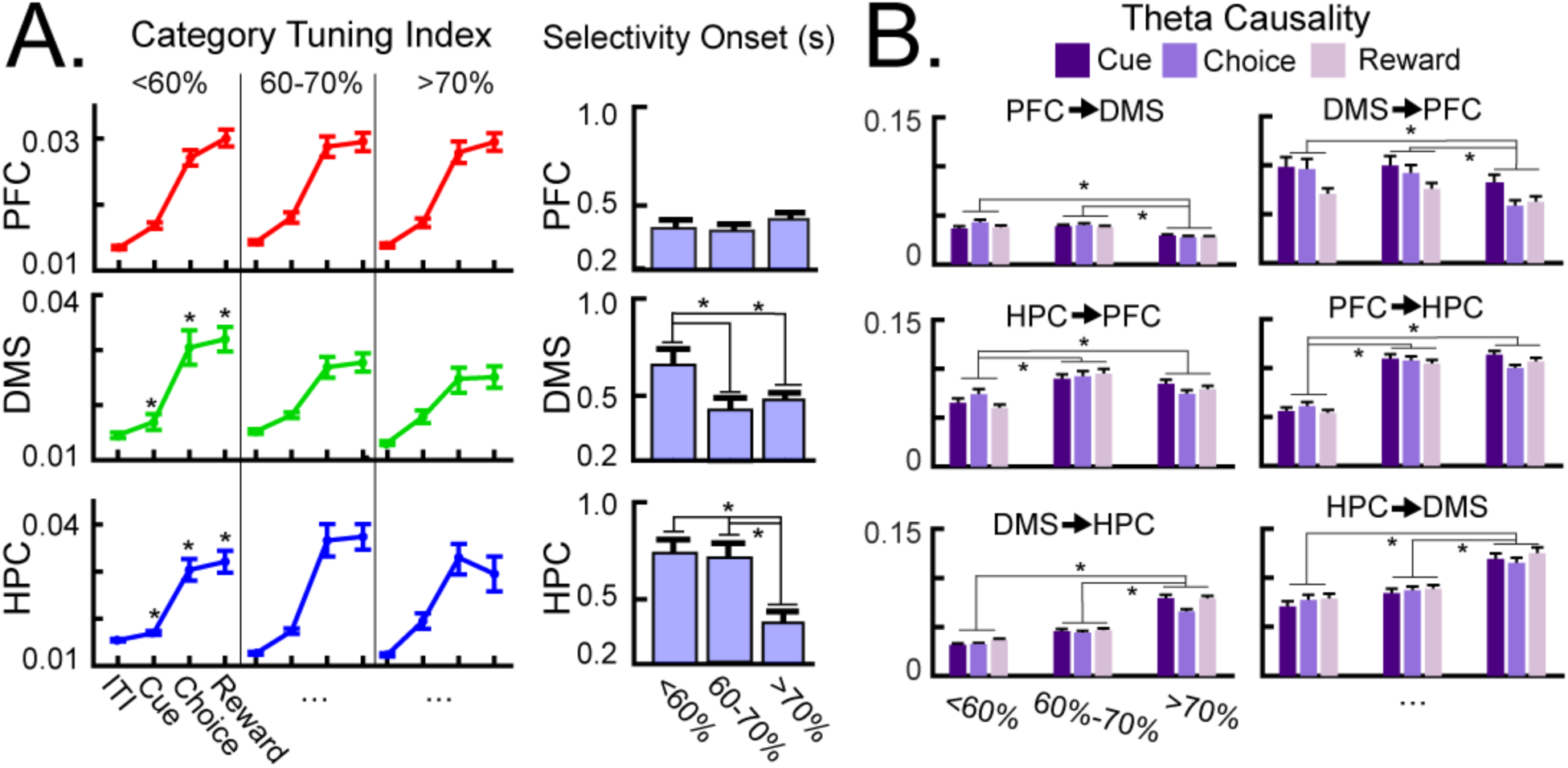
An analysis by session accuracy. The main measures in Figure 2D and Figure 6A were replotted according to session accuracy (i.e., <60%, 60%-70%, and >70%), rather than by training phase. The learning-related results were qualitatively equivalent between methods. **A,** Left: average CTI values separated by session accuracy. CTI during the Cue phase increased across accuracy levels in the DMS and HPC. CTI during the Choice phase increased with accuracy levels in the HPC and decreased with accuracy levels in the DMS. Right: selectivity onset separated by session accuracy. The latency of selectivity onset was faster in the PFC than the other regions. Latencies decreased across accuracy levels in the DMS and HPC. **B,** Average theta Granger causality separated by accuracy sessions. DMS→PFC theta causality decreased across accuracy levels, whereas HPC-PFC and DMS-HPC theta causality increased from the first accuracy level to the second and decreased from the second accuracy level to the third. All error bars indicate *S.E.M*.

## References

1. Shepard, R. N., Hovland, C. I., & Jenkins, H. M. (1961). Learning and memorization of classifications. Psychological Monographs: General and Applied, 75(13), 1–42. doi:10.1037/h0093825

2. Zeithamova, D., Mack, M. L., Braunlich, K., Davis, T., Seger, C. A., Van Kesteren, M. T., & Wutz, A. (2019). Brain mechanisms of concept learning. The Journal of Neuroscience, 39(42), 8259–8266. doi:10.1523/jneurosci.1166-19.2019

3. Seger, C. A., & Miller, E. K. (2010). Category learning in the brain. Annual Review of Neuroscience, 33(1), 203–219. doi:10.1146/annurev.neuro.051508.135546

4. Freedman, D. J., Riesenhuber, M., Poggio, T., & Miller, E. K. (2001). Categorical representation of visual stimuli in the primate prefrontal cortex. Science, 291(5502), 312–316. doi:10.1126/science.291.5502.312

5. Mack, M. L., Preston, A. R., & Love, B. C. (2020). Ventromedial prefrontal cortex compression during concept learning. Nature Communications, 11(1), 1–11. doi: 10.1038/s41467-019-13930-8

6. Broschard, M. B., Turner, B. M., Tranel, D., & Freeman, J. H. (2024). Dissociable roles of the dorsolateral and ventromedial prefrontal cortex in human categorization. The Journal of Neuroscience, 44(34). 10.1523/jneurosci.2343-23.2024

7. Mack, M. L., Love, B. C., & Preston, A. R. (2016). Dynamic updating of hippocampal object representations reflects new conceptual knowledge. Proceedings of the National Academy of Sciences, 113(46). doi:10.1101/071118

8. Theves, S., Fernández, G., & Doeller, C. F. (2020). The Hippocampus Maps Concept Space, Not Feature Space. The Journal of Neuroscience, 40(38), 7318–7325. 10.1523/jneurosci.0494-20.2020

9. Bowman, C.R., Zeithamova, D. (2018). Abstract memory representations in the ventromedial prefrontal cortex and hippocampus support concept generalization. Journal of Neuroscience, 38(10), 2605–2614.

10. Knowlton, B. J., Mangels, J. A., & Squire, L. R. (1996). A Neostriatal Habit Learning System in Humans. Science, 273(5280), 1399–1402. doi:10.1126/science.273.5280.1399

11. Antzoulatos, E., & Miller, E. (2011). Differences between neural activity in prefrontal cortex and striatum during learning of novel abstract categories. Neuron, 71(2), 243–249. doi:10.1016/j.neuron.2011.05.040

12. Frank, M. J. (2005). Dynamic dopamine modulation in the basal ganglia: a neurocomputational account of cognitive deficits in medicate and nonmedicated parkinsonism. Journal of Cognitive Neuroscience, 17(1), 51–72.

13. Humphries, M. D., Stewart, R. D., Gurney, K. N. (2006). A physiologically plausible model of action selection and oscillatory activity in the basal ganglia. Journal of Neuroscience, 26(1), 12921–12942.

14. Seger, C. A. (2008). How do the basal ganglia contribute to categorization? Their roles in generalization, response selection, and learning via feedback. Neuroscience & Biobehavioral Reviews, 32, 265–278.

15. Ashby, F. G., Alfonso-Reese, L. A., Turken, A. U., & Waldron, E. M. (1998). A neuropsychological theory of multiple systems in category learning. Psychological Review, 105(3), 442–481. 10.1037//0033-295X.105.3.442

16. Mack, M. L., Love, B. C., & Preston, A. R. (2018). Building concepts one episode at a time: The hippocampus and concept formation. Neuroscience Letters, 680, 31–38. doi:10.1016/j.neulet.2017.07.061

17. Love, B. C., & Gureckis, T. M. (2007). Models in search of a brain. *Cognitive, Affective*, & Behavioral Neuroscience, 7(2), 90–108.

18. Broschard, M. B., Kim, J., Love, B. C., Wasserman, E. A., & Freeman, J. H. (2021). Prelimbic cortex maintains attention to category-relevant information and flexibly updates category representations. Neurobiology of Learning and Memory, 185, 107524. doi:10.1016/j.nlm.2021.107524

19. Broschard, M. B., Kim, J., Love, B. C., Halverson, H. E., & Freeman, J. H. (2024). Disrupting dorsal hippocampus impairs category learning in rats. Neurobiology of Learning and Memory, 212, 10.1016/j.nlm.2024.107941.

20. Broschard, M. B., Kim, J., Love, B. C., & Freeman, J. H. (2023). Dorsomedial striatum, but not dorsolateral striatum, is necessary for rat category learning. Neurobiology of Learning and Memory, 199, 107732. 10.1016/j.nlm.2023.107732

21. Love, B. C., Medin, D. L., & Gureckis, T. M. (2004). Sustain: A network model of category learning. Psychological Review, 111(2), 309–332. doi:10.1037/0033-295x.111.2.309

22. Ashby, F. G., & Maddox, W. T. (2011). Human category learning 2.0: Human category learning 2.0. Annals of the New York Academy of Sciences, 1224(1), 147–161. 10.1111/j.1749-6632.2010.05874.x

23. Weichart, E. R., Evans, D. G., Galdo, M., Bahg, G., & Turner, B. M. (2022). Distributed Neural Systems support flexible attention updating during category learning. Journal of Cognitive Neuroscience, 34(10), 1761–1779. 10.1162/jocn_a_01882

24. Rehder, B., & Hoffman, A. B. (2005). Eyetracking and selective attention in category learning. Cognitive Psychology, 51(1), 1–41. 10.1016/j.cogpsych.2004.11.001

25. Gao, M., Turner, B. M., & Sloutsky, V. M. (2024). The role of attention in category representation. Cognitive Science, 48(4). 10.1111/cogs.13438

26. Braunlich, K., & Love, B. C. (2022). Bidirectional influences of information sampling and concept learning. Psychological Review, 129(2), 213–234. 10.1037/rev0000287

27. Kruschke, J. K. (2001). Toward a unified model of attention in associative learning. Journal of Mathematical Psychology, 45(6), 812–863. 10.1006/jmps.2000.1354

28. Price, A., Filoteo, J. V., & Maddox, W. T. (2009). Rule-based category learning in patients with parkinson’s disease. Neuropsychologia, 47(5), 1213–1226. 10.1016/j.neuropsychologia.2009.01.031

29. Andersen, S. K., & Müller, M. M. (2010). Behavioral performance follows the time course of neural facilitation and suppression during cued shifts of feature-selective attention. Proceedings of the National Academy of Sciences, 107(31), 13878–13882. 10.1073/pnas.1002436107

30. Bridwell, D. A., & Srinivasan, R. (2012). Distinct attention networks for feature enhancement and suppression in vision. Psychological Science, 23(10), 1151–1158. 10.1177/0956797612440099

31. Abdi, H., & Williams, L. J. (2010). Principal component analysis. WIREs Computational Statistics, 2(4), 433–459. 10.1002/wics.101

32. Lee, H., Battle, A., Raina, R., & Ng, A. Y. (2007). Efficient sparse coding algorithms. Advances in Neural Information Processing Systems 19, 801–808. 10.7551/mitpress/7503.003.0105

33. Duan, Y., Zhan, J., Gross, J., Ince, R. A. A., & Schyns, P. G. (2024). Pre-frontal cortex guides dimension-reducing transformations in the occipito-ventral pathway for categorization behaviors. Current Biology, 34(15). 10.1016/j.cub.2024.06.050

34. Ashby, F. G., & Rosedahl, L. (2017). A neural interpretation of exemplar theory. Psychological Review, 124(4), 472–482. 10.1037/rev0000064

35. Robbins, T. W., & Everitt, B. J. (1992). Functions of dopamine in the dorsal and ventral striatum. Seminars in Neuroscience, 4(2), 119–127. 10.1016/1044-5765(92)90010-y

36. Balleine, B. W., Peak, J., Matamales, M., Bertran-Gonzalez, J., & Hart, G. (2021). The dorsomedial striatum: An optimal cellular environment for encoding and updating goal-directed learning. Current Opinion in Behavioral Sciences, 41, 38–44. 10.1016/j.cobeha.2021.03.004

37. Mok, R. M., & Love, B. C. (2019). A non-spatial account of place and grid cells based on clustering models of concept learning. Nature Communications, 10(1). 10.1038/s41467-019-13760-8

38. Aronov, D., Nevers, R., & Tank, D. W. (2017). Mapping of a non-spatial dimension by the hippocampal–entorhinal circuit. Nature, 543(7647), 719–722. 10.1038/nature21692

39. Love, B. C., & Tomlinson, M. (2010). Mechanistic models of associative and rule-based category learning. The Making of Human Concepts, 53–74. 10.1093/acprof:oso/9780199549221.003.04

40. Zeithamova, D., Dominick, A. L., & Preston, A. R. (2012). Hippocampal and ventral medial prefrontal activation during retrieval-mediated learning supports novel inference. Neuron, 75(1), 168–179. 10.1016/j.neuron.2012.05.010

41. Backus, A. R., Schoffelen, J., Szebényi, S., Hanslmayr, S., & Doeller, C. F. (2016). Hippocampal-Prefrontal Theta Oscillations Support Memory Integration. Current Biology, 26(4), 450–457. doi:10.1016/j.cub.2015.12.048

42. Helie, S., Roeder, J. L., & Ashby, F. G. (2010). Evidence for cortical automaticity in rule-based categorization. The Journal of Neuroscience, 30(42), 14225–14234. 10.1523/jneurosci.2393-10.2010

43. Ashby, F. G., Ennis, J. M., & Spiering, B. J. (2007). A neurobiological theory of automaticity in perceptual categorization. Psychological Review, 114(3), 632–656. 10.1037/0033-295x.114.3.632

44. Buetfering, C., Zhang, Z., Pitsiani, M., Smallridge, J., Boven, E., McElligott, S., & Häusser, M. (2022). Behaviorally relevant decision coding in primary somatosensory cortex neurons. Nature Neuroscience, 25(9), 1225–1236. 10.1038/s41593-022-01151-0

45. Yartsev, M. M., Hanks, T. D., Yoon, A. M., & Brody, C. D. (2018). Causal contribution and dynamical encoding in the striatum during evidence accumulation. eLife, 7. 10.7554/elife.34929

46. Nieh, E. H., Schottdorf, M., Freeman, N. W., Low, R. J., Lewallen, S., Koay, S. A., Pinto, L., Gauthier, J. L., Brody, C. D., & Tank, D. W. (2021). Geometry of abstract learned knowledge in the hippocampus. Nature, 595(7865), 80–84. 10.1038/s41586-021-03652-7

47. Hanks, T. D., Kopec, C. D., Brunton, B. W., Duan, C. A., Erlich, J. C., & Brody, C. D. (2015). Distinct relationships of parietal and prefrontal cortices to evidence accumulation. Nature, 520(7546), 220–223. 10.1038/nature14066

48. Rigotti, M., Barak, O., Warden, M. R., Wang, X.-J., Daw, N. D., Miller, E. K., & Fusi, S. (2013). The importance of mixed selectivity in complex cognitive tasks. Nature, 497(7451), 585–590. 10.1038/nature12160

49. Parthasarathy, A., Herikstad, R., Bong, J. H., Medina, F. S., Libedinsky, C., & Yen, S.-C. (2017). Mixed selectivity morphs population codes in prefrontal cortex. Nature Neuroscience, 20(12), 1770–1779. 10.1038/s41593-017-0003-2

50. Benoit, R. G., Hulbert, J. C., Huddleston, E., & Anderson, M. C. (2015). Adaptive top–down suppression of hippocampal activity and the purging of intrusive memories from consciousness. Journal of Cognitive Neuroscience, 27(1), 96–111. doi:10.1162/jocn_a_00696

51. Topolnik, L., & Tamboli, S. (2022). The role of inhibitory circuits in hippocampal memory processing. Nature Reviews Neuroscience, 23(8), 476–492. 10.1038/s41583-022-00599-0

52. Jeong, N., & Singer, A. C. (2022). Learning from inhibition: Functional roles of hippocampal CA1 inhibition in spatial learning and memory. Current Opinion in Neurobiology, 76, 102604. 10.1016/j.conb.2022.102604

53. McKenzie, S. (2017). Inhibition shapes the organization of hippocampal representations. Hippocampus, 28(9), 659–671. 10.1002/hipo.22803

54. Chudasama, Y., Doobay, V. M., & Liu, Y. (2012). Hippocampal-prefrontal cortical circuit mediates inhibitory response control in the rat. The Journal of Neuroscience, 32(32), 10915–10924. 10.1523/jneurosci.1463-12.2012

55. Anderson, M. C., Bunce, J. G., & Barbas, H. (2017). Prefrontal–hippocampal pathways underlying inhibitory control over memory. Neurobiology of Learning and Memory, 134, 145–161. 10.1016/j.nlm.2015.11.008

56. Malik, R., Li, Y., Schamiloglu, S., & Sohal, V. S. (2022). Top-down control of hippocampal signal-to-noise by prefrontal long-range inhibition. Cell, 185(9). 10.1016/j.cell.2022.04.001

57. Church, B. A., Krauss, M. S., Lopata, C., Toomey, J. A., Thomeer, M. L., Coutinho, M. V., Volker, M. A., & Mercado, E. (2010). Atypical categorization in children with high-functioning autism spectrum disorder. Psychonomic Bulletin & Review, 17(6), 862–868. 10.3758/pbr.17.6.862

58. Zaki, S. R. (2004). Is categorization performance really intact in amnesia? A meta-analysis. Psychonomic Bulletin & Review, 11(6), 1048–1054. 10.3758/bf03196735

59. Tolman, E. C. (1948). Cognitive maps in rats and men. Psychological Review, 55(4), 189–208. 10.1037/h0061626

60. Manns, J. R., & Eichenbaum, H. (2009). A cognitive map for object memory in the hippocampus. Learning & Memory, 16(10), 616–624. 10.1101/lm.1484509

61. Gilboa, A., & Marlatte, H. (2017). Neurobiology of schemas and schema-mediated memory. Trends in Cognitive Sciences, 21(8), 618–631. 10.1016/j.tics.2017.04.013

62. Broschard, M. B., Kim, J., Love, B. C., & Freeman, J. H. (2020). Category learning in rodents using touchscreen-based tasks. *Genes*, Brain and Behavior. doi:10.1111/gbb.12665

63. Crijns, E., & Op de Beeck, H. (2019). The Visual Acuity of Rats in Touchscreen Setups. Vision, 4(1), 4. 10.3390/vision4010004

64. Broschard, M. B., Kim, J., Love, B. C., Wasserman, E. A., & Freeman, J. H. (2019). Selective attention in rat visual category learning. Learning & Memory, 26(3), 84–92. doi:10.1101/lm.048942.118

65. Paxinos, G. and Watson, C. (1998) The Rat Brain in Stereotaxic Coordinates. Academic Press, San Diego.

66. Hearst, M. A., Dumais, S. T., Osuna, E., Platt, J., & Scholkopf, B. (1998). Support Vector Machines. IEEE Intelligent Systems and Their Applications, 13(4), 18–28. 10.1109/5254.708428

67. King, J.-R., & Dehaene, S. (2014). Characterizing the dynamics of mental representations: The temporal generalization method. Trends in Cognitive Sciences, 18(4), 203–210. 10.1016/j.tics.2014.01.002

68. Shojaie, A., & Fox, E. B. (2022). Granger causality: A review and recent advances. Annual Review of Statistics and Its Application, 9(1), 289–319. 10.1146/annurev-statistics-040120-010930

69. Fiebelkorn, I. C., Pinsk, M. A., & Kastner, S. (2018). A Dynamic Interplay within the frontoparietal network underlies rhythmic spatial attention. Neuron, 99(4). 10.1016/j.neuron.2018.07.038

70. Seth, A. K., Barrett, A. B., & Barnett, L. (2015). Granger causality analysis in neuroscience and neuroimaging. The Journal of Neuroscience, 35(8), 3293–3297. 10.1523/jneurosci.4399-14.2015

